# SifiNet: A robust and accurate method to identify feature gene sets and annotate cells

**DOI:** 10.1101/2023.05.24.541352

**Authors:** Qi Gao, Zhicheng Ji, Liuyang Wang, Kouros Owzar, Qi-Jing Li, Cliburn Chan, Jichun Xie

## Abstract

SifiNet is a robust and accurate computational pipeline for identifying distinct gene sets, extracting and annotating cellular subpopulations, and elucidating intrinsic relationships among these subpopulations. Uniquely, SifiNet bypasses the cell clustering stage, commonly integrated into other cellular annotation pipelines, thereby circumventing potential inaccuracies in clustering that may compromise subsequent analyses. Consequently, SifiNet has demonstrated superior performance in multiple experimental datasets compared with other state-of-the-art methods. SifiNet can analyze both single-cell RNA and ATAC sequencing data, thereby rendering comprehensive multiomic cellular profiles. It is conveniently available as an open-source R package.

## Introduction

Single-cell sequencing techniques, including single-cell RNA sequencing (scRNA-seq) and single-cell ATAC sequencing (scATAC-seq), allow researchers to quantify cellular omic phenotypes. An ideal single-cell data analysis is expected to help researchers understand cellular omic heterogeneity, extract the cell subpopulations of interest, identify feature gene sets corresponding to cell subpopulations, as well as reveal the relationship between cell subpopulations.

Among these analysis tasks, identifying feature gene sets is a crucial step. Feature gene sets are defined as gene sets that are differentially expressed between cell subpopulations. They are often used to annotate cell subpopulations and to perform gene set enrichment analysis. Existing feature gene identification methods often employ a two-step approach (called two-step methods thereafter): cells are first clustered (such as Seurat[1–4], simple Louvain[5], Clustering through Imputation and Dimensionality Reduction (CIDR)[6], and SCANPY[7]) and differentially expressed gene (DEG) analysis (such as DESeq2[8], edgeR[9–13], limma[14,15], limma-voom[16], and MAST[17]) is then performed across cell clusters to identify cluster-specific feature genes. However, this approach has questionable accuracy on data with complex or subtle heterogeneity, since an inaccurate initial clustering step could lead to subsequent erroneous feature gene identification[18]. Alternatively, some methods identify feature genes by detecting highly variable genes (HVG) based on their deviations from the population with respect to deviance of model fitting[19], dropout rates[20], and UMI count distributions[21] (called HVG methods thereafter). However, these methods do not separate feature genes into subpopulation-specific gene sets, limiting their utility for annotating cells.

To overcome the limitations of existing methods, we present SifiNet, a unique approach for direct identification of feature gene sets that eliminates the need for prior cell clustering. Stemming from the crucial observation that genes co-differentially expressed within a cell subpopulation also exhibit co-expression patterns (Suppl. Note 1), SifiNet builds a gene co-expression network and inspects its topology to discern feature gene sets. These gene sets serve to compute cellular gene set enrichment scores and subsequently annotate cells. Additionally, networks among these gene sets imply relationships between cell subpopulations (Fig. 1). Furthermore, SifiNet can optionally integrate scATAC-seq data, as it forms a gene co-open-chromatin network and explores its topology to determine epigenomic feature gene sets. The ability of SifiNet to analyze both scRNA-seq and scATAC-seq data endows researchers with insight into cellular multiomic heterogeneity.

**Figure 1:**
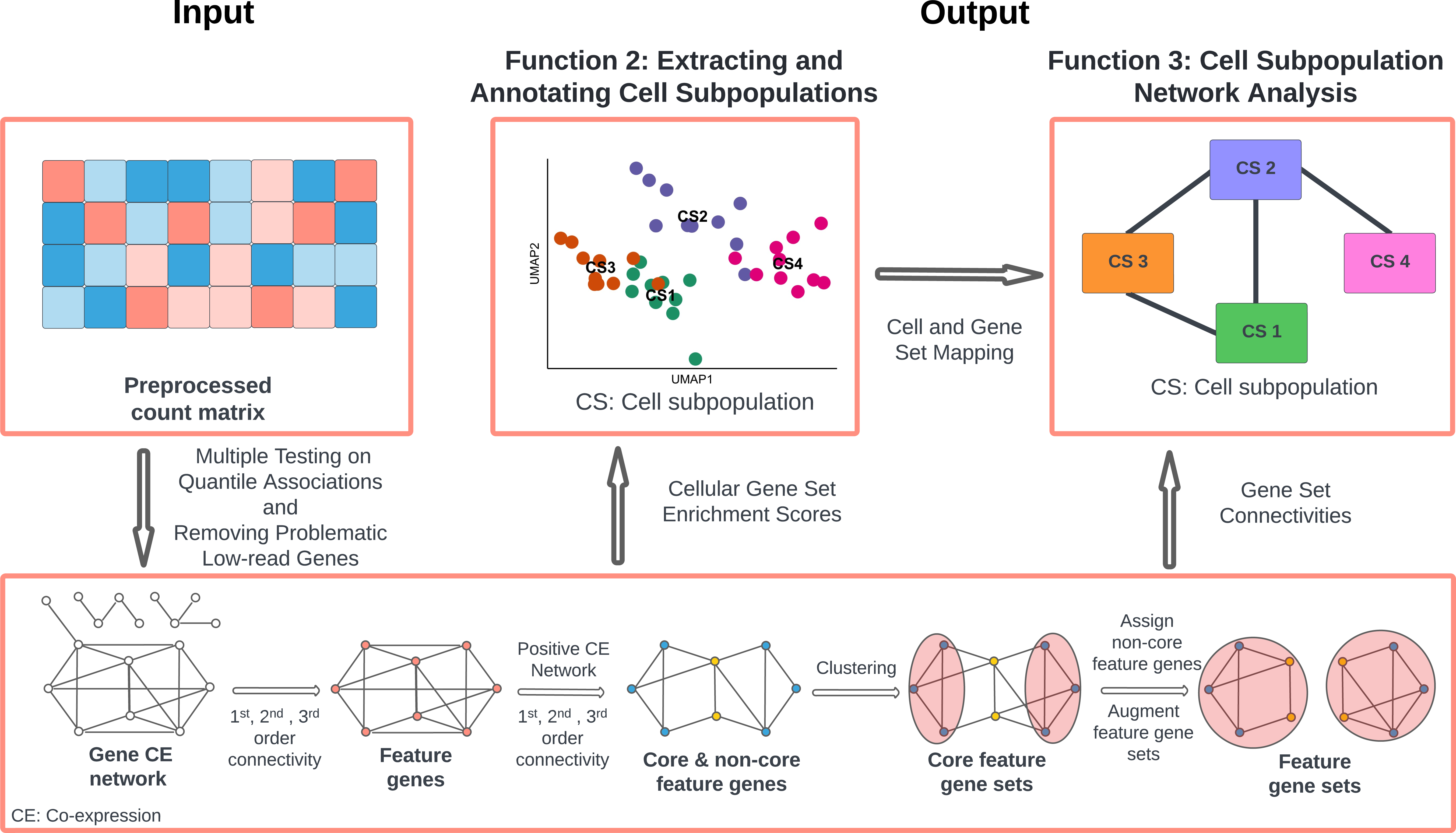
SifiNet pipeline. SifiNet takes the preprocessed feature count matrix as input, and uses gene co-expression network topology to identify feature gene sets (function 1). The identified feature gene sets are used to annotate cells (function 2), and the gene co-expression network is also used to reveal cell subpopulations’ transition or developmental relationships (function 3). The bottom row illustrates the main steps for identifying feature gene sets. After multiple testing on quantile associations and removing problematic low-read-count genes, SifiNet obtains a large gene co-expression network. Then, based on the 1^st^, 2^nd^, and 3^rd^ order connectivities, SifiNet identifies feature genes marked as red nodes. Then, SifiNet focuses on the positive co-expression network among feature genes and their node topologies to identify core feature genes, marked as blue nodes; the non-core features are marked in yellow. Next, SifiNet clustered the core feature genes into different clusters, and then assigned non-core feature genes and multi-role feature genes to the corresponding clusters. Finally, SifiNet obtains multiple feature gene sets.

We demonstrate that SifiNet outperforms existing two-step methods and HVG methods in identifying feature gene sets and enhancing cell annotation accuracy. Moreover, we illustrate that SifiNet can identify complex heterogeneity among cells and reveal potential developmental lineages among cell subpopulations. SifiNet can also be scaled to analyze datasets with millions of cells. We applied SifiNet to five published experimental datasets and uncovered several potentially novel discoveries, such as potential new cell cycle markers and senescence markers, the senescent-cell enriched subpopulation, the developmental lineage of myeloid progenitor cells, and the CD8 cell subpopulations along with their possible transition paths.

## Materials and Methods

### Computation Pipeline of SifiNet

SifiNet is a versatile pipeline with multiple functions, including the identification of feature gene sets, recognition of cell subpopulations, and exploration of potential developmental lineage among cell subpopulations (Fig. 1). The main innovative step is to use gene co-expression network topology to identify feature gene sets. Feature genes are genes over-expressed within a cell subpopulation; those under-expressed genes are treated as the feature genes of the remaining cells due to identifiability issues. A set of these genes, exhibiting co-over-expression within the same cell subpopulation, constitutes a feature gene set.

To identify feature gene sets, SifiNet first constructs a gene co-expression network based on quantile association measures. Then, based on topology measures, we choose the genes that form close communities. These genes are likely to be feature genes. Next, we focus on the positive co-expression network of those genes and their community patterns to cluster genes into multiple core feature gene sets. These core feature gene sets will be further enriched by adding additional genes, such as shared feature genes and genes performing multi-role functions. Based on the feature gene sets, we calculate the cellular gene set enrichment scores and annotate cells. Finally, we construct a cell subpopulation network to reveal the potential developmental or transitional relationship among cell subpopulations.

Let *Y_ic_* be the feature values of gene *i* in cell *c*. For scRNA-seq data, the gene feature value could be the gene UMI count; for scATAC-seq data, it could be the aggregated gene-level cut number. Based on the gene feature values, we construct a gene co-expression network among all the genes. Each node represents a gene, and each edge represents the co-expression between two genes. Let *s_c_* be cell *c*’s library size (such as total read counts). We use quantile associations to define gene co-expressions.

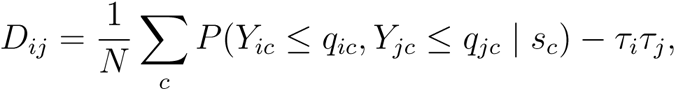

where *q_ic_* and *q_jc_* are the *τ_i_*-th and *τ_j_*-th conditional quantiles of gene *i* and gene *j*’s expressions. *s_c_* can be further extended to include other cell-specific covariates. Our previous study shows that quantile associations can effectively measure general dependence, especially for count data such as RNA-seq data[22]. In the gene-co-expression network, if *D_ij_* ≠ 0, there is an edge between nodes *i* and *j*.

Here, we establish a standardized estimator *S_ij_* for the co-expression *D_ij_*:

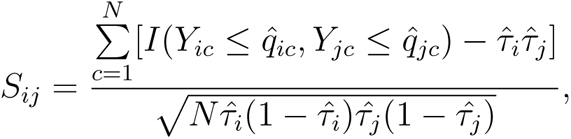

where 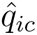 is the estimated cell specific *τ_i_*-th conditional quantile of gene *i*, and 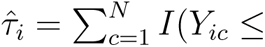 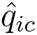)/*N*. By default, we use the quantile regression model *q_ic_* = *α_i_*_0_ + *α_i_*_1_*s_c_* to estimate 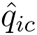, where 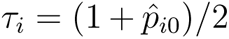 and 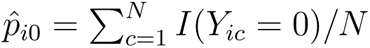 is the proportion of zero observations of gene *i*. In this default setting, the estimated 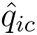 corresponds to the 50%-quantile among the non-zero observations of gene *i* given cell library size. However, when gene *i* has very low expression (when the median of *Y_ic_* is equal to 0 or *Y_ic_* are either 0 or 1), we set 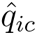 = 0 for stability and robustness consideration.

Assume *Y_ic_* and *Y_jc_* are independent given a specific cell *c*. Suppl. Note 1 shows that if gene *i* or gene *j* is not a feature gene, then *D_ij_* = 0, *i.e.*, these two genes are independent. Moreover, Suppl. Note 2 shows that under this case, the standardized estimator *S_ij_*asymptotically follows a standard normal distribution.

In reality, even given a cell, some genes may be dependent due to their biological interactions. For example, one gene may regulate another, causing two genes to co-express. Besides, hub genes usually co-express with many other genes. However, we found that feature genes, pathway genes, and hub genes exhibit different local topologies in the co-expression network (Suppl. Fig. 1). Specifically, feature genes tend to form close communities, while pathway genes and hub genes do not.

To capture the community patterns of feature genes, we introduce first-order, second-order, and third-order connectivities – three node topology measures. Denote the gene co-expression network *G* = *{V, E}* with node set *V* and edge set *E* . Here, *E* = *∪E_i_*, where *E_i_* is gene *i*’s edge set. For any set *S*, *|S|* is its cardinality.

- First-order connectivity measures gene *i*’s edge proportions, that is, the number of the observed edges over all possible edges: *C*_1_*_i_* = *|E_i_|/*(*p −* 1).
- Second-order connectivity measures the edge proportions among gene *i*’s neighbors: *C*_2_*_i_* = *|E ∩ {*(*j, k*) : *j, k ∈ N_i_}|/|N_i_|*(*|N_i_| −* 1), where *N_i_* = *{j* : (*i, j*) *∈ E_i_}* is gene *i*’s neighbor set.
- Third-order connectivity measures the edge proportions among gene *i*’s 2^nd^-order neighbors (*i.e.*, neighbors’ neighbors), denoted by *C*_3_*_i_*. Because some genes have many 2^nd^-order neighbors, leading to a long computation time for *C*_3_*_i_*, we employ a sampling strategy to improve the computation efficiency (Suppl. Note 3).

As we anticipate that feature genes will form communities with other feature genes, we define a gene as a feature gene if it has high *C*_2_*_i_* and *C*_3_*_i_* and moderate or high *C*_1_*_i_*. Genes with low *C*_1_*_i_*, *C*_2_*_i_* and *C*_3_*_i_* could be genes on the pathways or hub genes. Some feature genes may serve multi-roles as pathway genes and/or hub genes. These genes will have different topology structures from other feature genes. We will identify these multi-role feature genes in the end, after we identify other feature genes.

#### SifiNet function 1: Identifying feature gene sets

SifiNet rests upon a key observation: co-differentially-expressed genes show co-expression patterns, even if the expressions of those genes are conditionally independent given the cell types (Suppl. Note 1). Here we list the main steps to identify feature gene sets.

##### Step 1: Initiating a gene co-expression graph

Suppose after preprocessing, the single-cell dataset contains *p* genes and *N* cells. Initiate gene co-expression graph by *G* = *{V, E}* with *V* = [*p*]. To obtain *E*, we set up *p*(*p −* 1)*/*2 hypotheses:

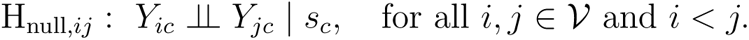

Then we perform multiple testing on H_null_*_,ij_* to determine the edges with FDR controlled at 5% (Suppl. Note 3).

##### Step 2: Eliminating problematic low-count genes and corresponding edges

Some low-count genes exhibit spurious co-expressions with many other low-count genes, and these low-count genes form a spurious community. These genes are removed from the gene co-expression network (Suppl. Note 3).

##### Step 3. Identifying feature genes

The feature gene set *V*_+_ contains the genes with *C*_1_*_i_ ≥* 5*/*(*p −* 1), *C*_2_*_i_ ≥* 0.4 and *C*_3_*_i_ ≥* 0.3 (Suppl. Note 3).

##### Step 4. Identifying core feature genes

Core feature genes are those unique feature genes of a cell subpopulation, often showing strong positive co-expressions (Suppl. Note 1). To find core feature genes, we first define a positive co-expression network *G*_+_ = *{V*_+_*, E*_+_*}*, where *E*_+_ contains the positive co-expression edges *E*_+_ = *{*(*i, j*) : (*i, j*) *∈ E, i, j ∈ V*_+_*, S_ij_ >* 0*}*. The 1^st^-, 2^nd^-, and 3^rd^-order connectivities for *G*_+_, denoted by *C*_+_,_1_*_i_*, *C*_+_,_2_*_i_*, and *C*_+_,_3_*_i_*, are calculated correspondingly. By default, the core feature genes are those with *C*_+_,_1_*_i_ ≥* 5*/*(*|V*_+_*| −* 1), *C*_+_,_2_*_i_ ≥* 0.7, and *C*_+_,_3_*_i_ ≥* 0.5. See Suppl. Fig. 2 for an illustration.

##### Step 5. Clustering core feature genes into gene sets

Taking the positive co-expression network restricted to the core feature genes *G*_+,core_, we apply the Louvain algorithm to cluster these genes into *K* gene sets, denoted by *V_k_*, where *k ∈* [*K*].

##### Step 6. Assigning non-core feature genes

A non-core feature gene may be shared by multiple feature gene sets. It is added to *V_k_* if it connects to more than 40% of the genes in *V_k_*.

##### Step 7. Augmenting feature gene sets with additional genes

Multi-role genes will have different connectivities from the sole feature genes. Thus, they might not be identified as feature genes in Step 3. For any gene not included in the gene sets, it will be added to *V_k_* if it connects more than 90% of the feature genes in *V_k_*.

Some steps involve setting tuning parameters, such as the 1^st^-, 2^nd^-, and 3^nd^-order connectivity thresholds, and the assignment criteria for non-core feature genes, etc. The default values of these parameters are set based on our experiences in analyzing multiple experimental datasets. In Step 5, the Louvain algorithm is applied with the default resolution parameter. Because the core feature genes usually exhibit a clear community structure, different resolution parameters usually lead to the same cluster results.

#### SifiNet function 2: Annotating cells with feature gene sets

Based on *V_k_*, *k ∈* [*K*], we calculate the cellular gene set enrichment scores *α_c,_*_1_*, . . . , α_c,K_* for any cell *c*. For all the data analyses in this paper, we used the gene set variation analysis (GSVA)[23] scores. We define the cell subpopulation corresponding to *V_k_* by

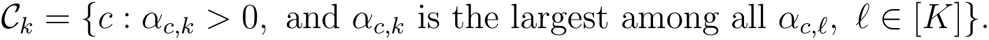

Those cells with all negative *α_c,k_* will be assigned to *C_K_*_+1_, called the unassigned group. These cells could be in the transitional stage or simply outliers. To annotate these unassigned cells, we can further apply DEG analysis to find their corresponding feature gene set.

If the feature gene set contains many matching gene markers from known marker gene databases, we can annotate the cell subpopulation with the corresponding cell type. See the Monoclonal dataset analysis for an example. Otherwise, we refer to this cell subpopulation as new, and annotate them with the identified feature genes. See Discussions for more details.

To visualize the cell subpopulations, we derive the GSVA score matrix for all the cells and all the gene sets. Then we calculate the first two principal components (PCs) of the GSVA score matrix. The first two PCs are used to plot the cell subpopulations in a 2D space. See Figs. 3d, 3e, 4e, 4f for examples.

For data with complex heterogeneity structure, after we cluster cells based on the gene set enrichment scores, SifiNet can be applied again to a subset of cells to explore their local heterogeneity structure. By sequentially applying SifiNet, we can identify the hierarchical structure of cell subpopulations. See the BoneMarrow dataset analysis for an example.

#### SifiNet function 3: Constructing cell subpopulation network

By using the *K* feature gene sets as nodes, we can construct a gene-set-level graph representing the cell subpopulation lineage. There are a total of *K*(*K −* 1)*/*2 possible edges. For edge (*k, ℓ*), its width is defined as edge proportions among all possible edges.

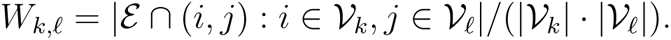

The width indicates the relationship between the two corresponding gene sets. Finally, the gene set network can be transferred to cell subpopulation network after identifying the cell subpopulation corresponding to each feature gene set. Then, a wide edge between two cell subpopulations suggests possible transitional or developmental relationships between them. By default, edges with width less than 0.3 will be removed from the visualization figure.

### Competing methods

We compared the performance of SifiNet against 5 two-step methods, 3 HVG identification methods, and 1 trajectory analysis method. See Suppl. Table 1 for details of the competing methods.

The two-step methods first use clustering methods to identify cell subpopulations, and then apply DEG analysis to identify cluster-specific DEGs. They are named after the DEG methods (including DESeq2[8], edgeR[9–13], limma[14,15], limma-voom[16], and MAST[17,24]) and the clustering methods (including Seurat[1–4], simple Louvain[5], and CIDR[6]), followed by TS. For example, DESeq2-Seurat-TS means that we first apply Seurat to cluster cells, and then use DESeq2 to identify cluster-specific DEGs. We tried all cross-over combinations of DEG and the clustering methods on SD1. CIDR utilizes both gene expression levels and dropout rates to cluster cells. It achieves the higher clustering performance in terms of adjusted rand index (ARI) for SD1 (Suppl. Fig. 3). Also, combined with DEG methods, CIDR achieves higher F1 score in feature gene identification compared with other clustering methods (Suppl. Fig. 4). Thus, we use CIDR as the default clustering method for all other analyses.

The clustering methods are applied at default resolution. The DEG methods are applied in similar ways as described in[25]. False discovery rate (FDR) is controlled at 0.05 with Benjamini-Hochberg (BH) procedure if there are p-values available in the method.

The oracle-cluster methods were also introduced to explain why SifiNet work better than the two-step methods. The oracle-cluster methods assume the ground truth of cell types, and then apply DEG analysis to identify cell-type-specific feature genes. The oracle-cluster methods include DESeq2-OC, edgeR-OC, limma-OC, limma-voom-OC, and MAST-OC.

The HVG identification methods identify feature genes through their deviation from the average gene properties. The HVG identification methods do not rely on knowing cell types or clustering. The methods we included are scry[26], M3Drop[27], and singleCellHaystack[21]. Because scry does not provide gene p-values, when we apply it to the simulated datasets, we selected the top *p*_1_ genes where *p*_1_ is the number of the actual feature genes.

Moreover, we also compared SifiNet with Monocle3[28–33] in their performance to construct cell subpopulation networks. Monocle3 is a comprehensive pipeline with a function to construct graph-based single-cell trajectories.

### Data Sets and Analysis

To benchmark SifiNet’s performance, we applied SifiNet to simulated and experimental datasets, including two simulated datasets (SD1, SD2) and five experimental datasets (Monoclonal[34], IPF1[35], BoneMarrow[36], CD8[37], MassiveRNA[38]). An additional experimental dataset IPF2[35] was used for validation. Suppl. Table 2 lists the details of the datasets.

#### Simulated Data

We used the R package GeneScape[39] to generate two numerical dataset, SD1 and SD2. GeneScape can simulate scRNA-seq data with customized differential expression structures, such as core and shared feature genes, sub-cell types, pathways and hub genes. The parameters used for simulation are shown in https://github.com/jichunxie/SifiNet_manu_ support.

SD1 simulated 100 repetitions of scRNA-seq datasets with three cell types; each dataset mimics the case where a rare cell subpopulation recently developed from another non-rare cell subpopulation. SD2 simulated five equal-sized cell subpopulations. Among them, two are transitional, and the other three are ending. Transitional cell types share feature genes with their neighboring ending cell subpopulations. The cell developmental lineage has the structure: ending-transition-ending-transition-ending. See Suppl. Note 4 for detailed simulation settings. For preprocessing, for all simulated datasets, we removed the cells with less than 10% genes expressed and genes expressed in less than 10% cells.

#### Experimental Data

##### Monoclonal

The dataset was collected from a monoclonal murine colorectal carcinoma cell line CT26WT. All cells were cloned from a single ancestral cell. We preprocessed the data in the following way. We first filter the genes similar as in the original study[34]. We then removed the cells whose mitochondrial gene expression takes more than 5% of the total gene expression, the cells whose total number of read count is greater than 1.8 times of the mean expression levels, the cells with less than 10% of the genes expressed, and the genes expressed in less than 10% of the cells. No additional normalization was performed before we applied SifiNet. This is because SifiNet constructed a gene co-expression graph based on quantile associations. The quantile regressions used to estimate quantile associations serve a similar role as quantile normalization in this step.

To curate a benchmark gene set to verify the identified genes, we did the following steps. First, we collected human cell cycle related genes using 3 libraries, Reactome[40,41], Bio-carta[42], KEGG[43–45], and 32 datasets from single studies or pathway analyses (Suppl. Table 3) with “CELLCYCLE” or “CELL CYCLE” in their name from GSEA MSigDB C2 curated human gene sets v2023.1[46–48]. Human genes that were included in the related pathway in at least 1 library or at least 3 studies were selected and converted to their mouse equivalents using biomaRt R package[49,50]. Second, for the Uniprot database, we downloaded the Uniprot gene annotations on May 1, 2023, and selected those genes whose “Biological process” annotations[51,52] include at least 1 of the following keywords: mitotic, centriole, centrosome, centromere, spindle, chromosome separation, cohesin, DNA replication, DNA repair, microtubule, nuclear chromosome segregation, cell cycle, cell division, cell growth. Finally, the intersection of selected genes in these two databases was chosen as the benchmark cell cycle set (475 genes). Among them, 437 genes remained in the Monoclonal data after preprocessing.

##### IPF1 and IPF2

These two datasets were collected from the existing study on idiopathic pulmonary fibrosis (IPF)[35]. IPF1 contains the pre-annotated “Ciliate” cells in IPF patients. IPF2 contains the pre-annotated “KRT5-/KRT17+” epithelial cells and the “Proliferating Epithelial Cells”. For preprocessing, we removed the cells with less than 5% genes expressed, and the genes expressed in less than 5% cells. No additional normalization was performed before we applied SifiNet.

Next, only for DEG analysis, cell counts were normalized by sctransform[53] in Seurat and the log fold changes and p-values are calculated using Seurat FindMarkers[8,24,29,54].

##### BoneMarrow

BoneMarrow was collected from an existing study on myeloid differentiation[36]. We focus on the pre-annotated “Unsorted myeloid” cells. Because BoneMarrow contains multiple batches of single-cell data, we used sctransform to regress out the batch effect by including a batch variable. Then, we removed the cells with fewer than 500 total UMI counts, and the genes expressed in fewer than 2% of the cells.

##### CD8

CD8 is a subset of a 10x scRNA-seq and scATAC-seq multiome datasets[37]. An annotation of cell types is provided by Seurat. CD8 dataset contains cells with “CD8 Näıve”, “CD8 TEM 1” and “CD8 TEM 2”. In scATAC-seq data, peaks were aggregated to represent the epigenomic features of the corresponding gene if they are located within the 5000-base-pair upstream of the transcription starting site (TSS) of gene. Only the peaks annotated by one gene are considered. Only genes available in both scRNA-seq data and scATAC-seq data were kept. In scRNA-seq data, cells with total read count greater than 1.8 times of the mean total read count were removed. In scATAC-seq data, cells with total epigenomic feature count greater than 1.8 times of the mean total epigenomic feature count were removed. We further removed cells with fewer than 2% of the genes detected in either scRNA-seq or scATAC-seq, and then the genes expressed in fewer than 2% of the cells in either scRNA-seq or scATAC-seq. No additional normalization was performed before we applied SifiNet.

##### MassiveRNA

MassiveRNA is a large dataset with more than 1 million cells[38]. We removed those cells with fewer than 2% of the genes expressed, and then those genes expressed in fewer than 2% of the cells. After that, 1287071 cells and 10448 genes remain in the dataset. SifiNet was applied to the whole dataset and downsampled datasets with number of cells 10000, 20000, 50000, 100000, 200000, 400000, 600000, 800000, and 1000000. No additional normalization was performed before we applied SifiNet. The time consumption was evaluated using the total elapsed time when running SifiNet on the dataset, and the peak memory usage was evaluated using the gc function in R.

## Results

Because SifiNet uses intrinsic gene co-expression network topology to identify feature gene sets, it is not affected by the clustering accuracy. Thus, SifiNet can effectively identify feature gene sets even when the cell subpopulation has complex heterogeneity structures and cannot be clustered well. For instance, some cell subpopulations may be transitioning to another state, some may share feature genes, and some may exhibit hierarchical structures. SifiNet can effectively handle these complex cases. By exploring multiple simulated and experimental datasets, we show that SifiNet has significantly improved accuracy in identifying feature gene sets and annotating cells compared to the competing methods, especially when cells display complex heterogeneity structures.

### SifiNet identifies feature gene sets with high accuracy

We applied SifiNet to SD1, Monoclonal, and IPF1, to identify the feature gene sets. SifiNet achieved a high accuracy in identifying known feature gene sets, and also identified potential new cell cycle markers and new ciliated cell senescence markers.

First, for the 100 replicated datasets in SD1, SifiNet achieves the highest median F1 score among all methods, almost doubling the second best (Fig. 2a). Among the competing methods, singleCellHaystack, limma-CIDR-TS, limma-voom-CIDR-TS, and MAST-CIDR-TS exhibit higher median sensitivity than SifiNet, but significantly lower median specificity. High specificity is crucial in feature gene identification studies due to the large number of candidate genes. For instance, each replication of SD1 dataset contains, on average, 83 feature genes and 3318 non-feature genes. A specificity as high as 0.9 will falsely label, on average, 332 non-feature genes as feature genes, substantially obscuring the signal of the true feature genes. SifiNet’s main advantage is its high specificity (close to 1), indicating that most identified genes are genuine feature genes.

**Figure 2:**
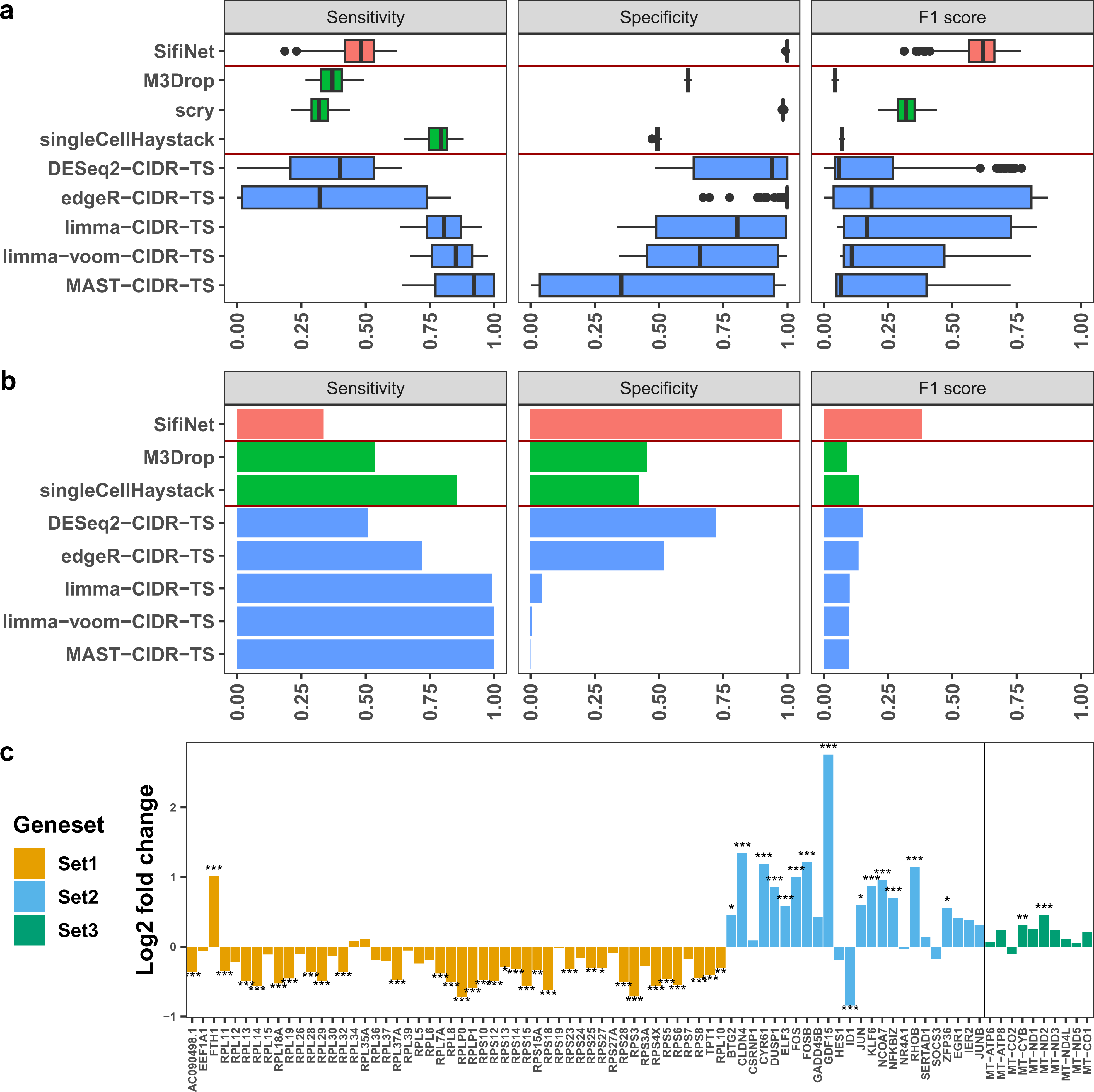
(a)-(b) Performance comparison between SifiNet and competing methods on SD1 and Monoclonal. Sensitivity, specificity, and F1 accuracy scores of 3 types of methods: SifiNet (red), HVG methods (green), and two-step methods (blue). (a) Performance on SD1, a simulated dataset with 100 replications. (b) Performance on Monoclonal to identify cell cycle marker genes. The benchmark cell cycle marker genes were curated from existing literature (Methods). **(c) Verification of SifiNet-identified feature genes with an independent dataset.** SifiNet identified the feature genes in IPF1. Then, DEG analysis was performed on IPF2 to show their log2 fold changes between KRT5-/KRT17+ (senescence-enriched) epithelial cells and proliferating (non-senescent) epithelial cells in IPF2. Stars represent the significance level of the log2 fold change, *: adjusted p-value *<*= 10*^−^*^2^, **: adjusted p-value *<*= 10*^−^*^4^, ***: adjusted p-value *<*= 10*^−^*^6^.

We further investigated why the commonly used two-step methods underperform on SD1. We compared them with the oracle methods, which assume the true cell type labels are known. All oracle-cluster methods perform well on SD1, exhibiting high sensitivity and specificity (Suppl. Fig. 4). This indicates that the key factor is the clustering accuracy. If the resulting clustering is perfect, the DEG step can effectively identify feature genes. However, in practice, exact cell types are unknown, and clustering methods can only provide a reasonable estimate. When these estimates are inaccurate, two-step methods perform poorly (Suppl. Fig. 5).

Second, we applied SifiNet to the Monoclonal dataset to identify the feature genes related to cell cycles. Because all cells were cloned from a single ancestral cell, we reasoned that the major cellular transcriptomic difference stems from their cell cycle difference. Based on the gene co-expression network topology (Suppl. Fig. 6), SifiNet identified two feature gene sets. Set 1 contains many late G1 and S (G1/S) phase genes, such as GINS, MCM, DNA polymerase genes, and RPA complex genes. Interestingly, it also includes HAUS genes (*Haus3*, *Haus5*, *Haus6*, *Haus8* ). Previously, HAUS genes were thought to be related to the centrosome cycle by maintaining the integrity of the centrosome. Although many other centrosome-associated genes are considered to be markers for the G2/M phase[55], studies[56,57] indicate that centrosome integrity might also be a checkpoint for the G1 to S phase transition. Without HAUS proteins maintaining the centrosome integrity, the cells may be unable to progress to the late G1/S phase. SifiNet analysis pinpointed that the differential expression of HAUS genes is strongly correlated with other late G1/S phase feature genes. Set 2 features are composed of many well-known G2 and M (G2/M) phase genes, including Tubulin and SKA. The full list of identified genes is in Suppl. Table 4.

We compared the performance of SifiNet and the two-step methods on Monoclonal. Using a benchmark cell cycle marker gene set collected from the MSigDB[46–48] and Uniprot[58] databases, we calculated the sensitivity, specificity, and F1 score of each method. SifiNet has the highest F1 score among all methods (Fig. 2b). The second best, CIDR-DESeq2-TS, has an F1 score lower than half of SifiNet’s. This is because CIDR-DESeq2-TS has identified too many irrelevant genes, leading to low specificity. In contrast, a higher proportion of SifiNet-identified feature genes overlap with the benchmark gene set, leading to high specificity (Suppl. Fig. 7).

Third, we applied SifiNet to IPF1 to identify senescence-related feature gene sets. IPF1 contains 10,375 ciliated cells from Idiopathic Pulmonary Fibrosis (IPF) human lung tissues. Prior research has demonstrated a close relationship between senescence and IPF[59,60]. Therefore, by exploring cell-type-specific feature genes, we expect to identify feature genes associated with senescence. SifiNet identified three feature gene sets (Suppl. Table 4). Set 1 includes numerous ribosomal protein genes, known to accumulate in senescent cells[61]. Set 2 contains *Jun*, *Junb*, *Fos*, and *Fosb* from the Activator Protein-1 (AP-1) transcription factor (TF) family. Other genes in Set 2, such as *Gdf15*, *Egr1*, and *Zfp36*, have been previously found to interact with the AP-1 TF family[62–64]. This family is known to be involved in proliferation, differentiation, and tumorigenesis[65], highly related to the senescence process[66]. Set 3 genes are all mitochondrial genes. Previous studies have shown that mitochondrial genes are related to the senescence-associated secretory phenotype (SASP)[67], and mitochondrial dysfunction is associated with senescence[68]. These three gene sets may relate to different aspects of senescence.

To verify the identified senescence genes, we checked their log fold change and significance levels using an independent dataset, IPF2 (Fig. 2c). IPF2 contains two cell subpopulations: KRT5-/KRT17+ epithelial cells and proliferating epithelial cells; the former are treated as senescent cells, and the latter non-senescent. Among the 84 SifiNet identified genes, 45 are significantly differentially expressed (adjusted p-values *<* 0.0001, Fig. 2c) between two cell subpopulations. The analysis of independent dataset IPF2 verified the quality of the identified feature genes.

### SifiNet identifies cell subpopulations with high accuracy

Here, we aim to identify cell subpopulations corresponding to different cell states, such as different cell cycle phases and senescence levels. Cells with the same functional cell type but different states only have subtle differences; thus, existing clustering methods often fail to separate them well. However, SifiNet achieves high accuracy in calling these cell subpopulations with subtle differences.

First, we applied SifiNet to Monoclonal to call cells in different cell cycles. Cells with high enrichment scores of feature gene sets 1 and 2 were annotated as late G1/S phase and G2/M phase, respectively, by mapping the feature genes in the gene set to known cell cycle-related markers (Suppl. Table 4); the unassigned cells are annotated as early G1 phase. Fig. 3a displays the Uniform Manifold Approximation and Projection (UMAP)[28] plot of the cells annotated by SifiNet. SifiNet identified gene sets show strong within-set connectivity and relatively weaker between-set connectivity (Suppl. Fig. 6d). To set up a benchmark, we also calculated the Seurat cell cycle scores using known cell cycle markers[4,69] (Fig. 3b). Similar to SifiNet-identified feature genes, these two known cell cycle gene sets exhibit high within-set connectivity and relatively weaker between-set connectivity (Suppl. Fig. 6e). Although SifiNet does not utilize any known marker information, its annotation closely resembles the benchmark annotation (Suppl. Fig. 8). As a comparison, we applied CIDR to cluster cells into three cell clusters. The clustering results are less satisfactory: notably, CIDR clustered many early G1 and G2/M phase cells to the late G1/S phase (Fig. 3c). Overall, CIDR has a lower ARI score than SifiNet when compared with the benchmark (Fig. 3c, Suppl. Fig. 8). Furthermore, CIDR, with the default resolution parameter, yielded 8 clusters for cell clustering; the results were hard to interpret. (Suppl. Fig. 9).

**Figure 3:**
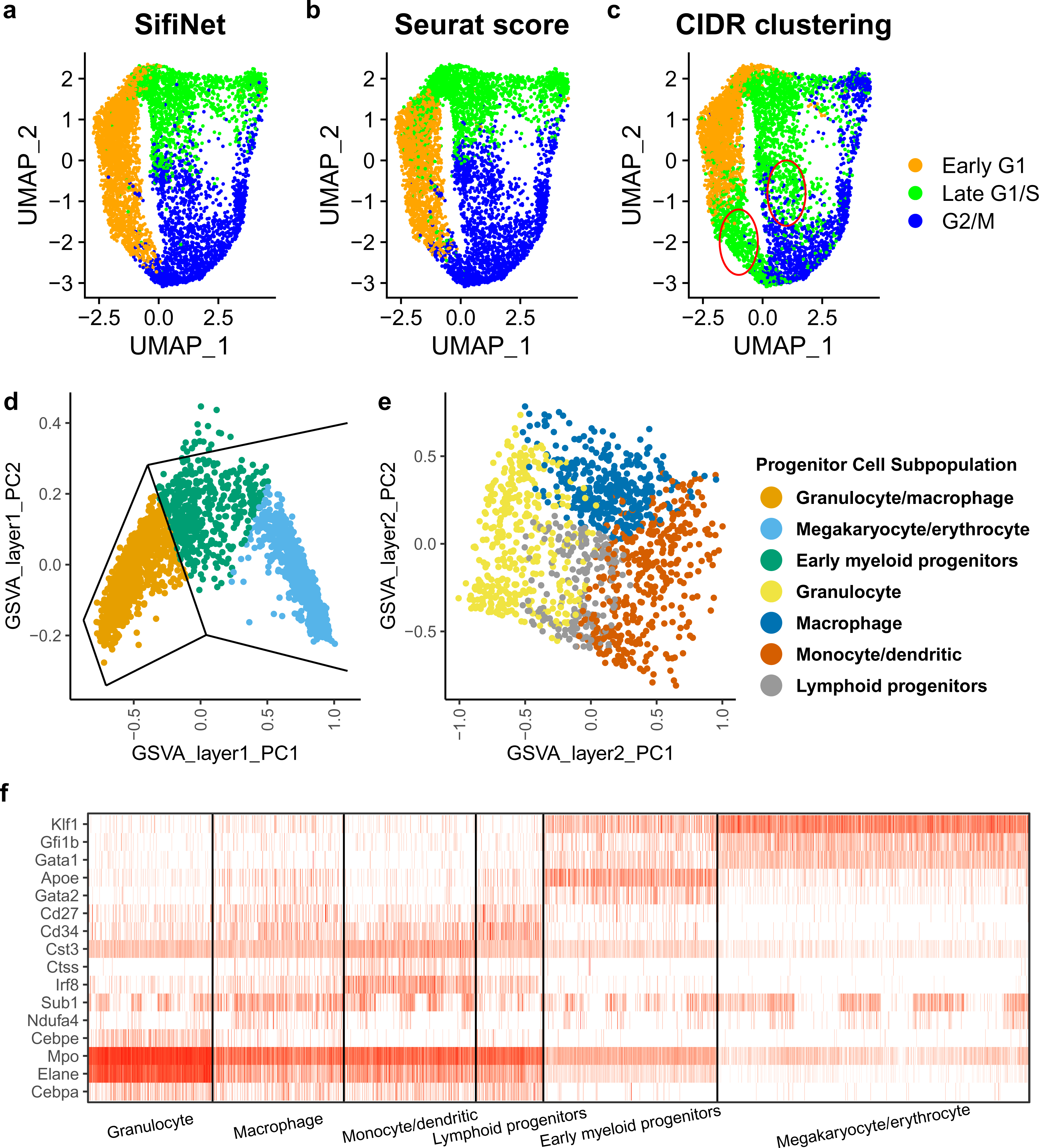
(a)-(c) UMAPs of Monoclonal cells. The cells were annotated by (a) SifiNet or (b) Seurat cell cycle scores, or clustered by (c) CIDR. For CIDR, the cell clusters are annotated to best map to the Seurat cell cycle score annotation. Red circles mark the major difference between the CIDR annotation and the benchmark annotation. **(d)-(f) SifiNet-annotated BoneMarrow cell subpopulations.** The x- and y-axis of (d) and (e) are the first two principal components (PCs) of the GSVA scores based on the SifiNet-identified gene sets. (d) When applied to all cells, SifiNet annotated three cell subpopulations: early progenitors, megakaryocyte/erythrocyte progenitor cells, and granulocyte/macrophage progenitor cells. (e) Applied to the granulocyte/macrophage progenitor cells, SifiNet identified four cell subpopulations: Granulocyte progenitor cells, Macrophage progenitor cells, Monocyte and dendritic progenitor cells, and T and B progenitor cells. T and B cells are not myeloid cells, but all these cells share common progenitor cells, hematopoietic stem cells. These cells may get mixed up with the myeloid progenitor cells. (f) Heatmap of some marker gene expressions in the annotated cell subpopulations.

Second, we applied the SifiNet to label the senescent cells in IPF1. To call the SifiNet-identified senescent cells in IPF1, we calculated the GSVA scores of the three gene sets. The 594 cells with top/bottom 40% GSVA scores of all three gene sets are labeled as SifiNet-senescent cells: top scores were used for up-regulated gene set (Gene Set 2), and bottom scores were used for down-regulated gene sets (Gene sets 1 and 3). Then, we check whether the widely known senescence marker, *Cdkn1a* [70], is differentially expressed in the SifiNet-senescent cells versus the others. It turns out that *Cdkn1a* has log2 fold change 0.636, and adjusted p-value= 1.54 *×* 10*^−^*^34^. As a comparison, we used SenMayo, a recently published senescence marker set[71], to label the senescence cells. We chose 594 cells (same number of cells as SifiNet-senescent cells) with the top SenMayo-GSVA scores. We found that *Cdkn1a* exhibited lower differential expression in the SenMayo-senescent cells (log2 fold change 0.437, adjusted p-value= 6.66 *×* 10*^−^*^9^). This suggests that SifiNet-identified gene sets annotated senescent ciliated cells better than the SenMayo senescence marker set.

### SifiNet revealed cellular hierarchical heterogeneity

Cellular heterogeneity is often hierarchical, especially for stem-like cells. We applied SifiNet to BoneMarrow, single-cell transcriptome dataset with heterogenous cells at various development stages isolated from mouse bone marrow. And SifiNet successfully predicted the developmental lineage of the myeloid progenitor cells before they grow into the downstream cells.

Taking the gene expression data from the whole cell set, SifiNet identified two feature gene sets in BoneMarrow, S1 and S2 (Suppl. Table 4) and separated the cells into three major populations with distinct transcriptomic features: early myeloid progenitor cells, megakaryocyte/erythrocyte progenitor cells, and granulocyte/macrophage progenitor cells (Fig. 3d). S1 contains markers for lineages in the granulocyte/macrophage progenitors[72,73], such as *Cebpa*. S2 contains markers for megakaryocyte/erythrocyte progenitors[74–76], such as *Gata1*, *Gfi1b*, and *Klf1* (Fig. 3f). The cells with positive S1 (S2) GSVA scores and negative S2 (S1) GSVA scores were clustered into granulocyte/macrophage progenitor cells (megakaryocyte/erythrocyte progenitor cells); those cells with both negative S1 and S2 GSVA scores were clustered into early progenitor cells. Early progenitor cell markers such as *Apoe* and *Gata2* [36] are significantly highly expressed in the early progenitor cells (*Apoe*: log2 fold change 1.75, adjusted p-value= 1.19 *×* 10*^−^*^169^; *Gata2* : log2 fold change 0.887, adjusted p-value= 2.77 *×* 10*^−^*^59^).

When applied to the annotated granulocyte/macrophage progenitor cells, SifiNet identified 3 feature gene sets S1a-S1c (Suppl. Table 4), dividing the cells into 4 subpopulations: granulocyte progenitor cells (corresponding to S1a), macrophage progenitor cells (corresponding to S1b), monocyte and dendritic progenitor cells (corresponding to S1c), and unassigned (Fig. 3e). DEG analysis shows that a marker gene for hematopoietic stem cells with the capacity to differentiate into T cells, *Cd34* [77], and memory B cell and T cell marker gene, *Cd27* [78,79], are significantly over-expressed in the unassigned subpopulation. Thus, we label this subpopulation as lymphoid progenitor cells. These cells may be mixed up with the myeloid progenitor cells when the original study sorted the cells[36].

### SifiNet revealed accurate cell subpopulation network in SD2

We applied SifiNet to SD2, a simulated dataset with five cell subpopulations representing a network with five lineage stages (Fig. 4b). While all the cells are distinctly separated into different clusters, the neighboring cell subpopulations share certain feature genes, indicating a developmental transition at the feature gene level (Suppl. Note 4). SifiNet identified five gene sets, separated the cells into five corresponding subpopulations, and constructed the network. Clearly, the cell subpopulation network built by SifiNet (Fig. 4a) is identical to the actual network (Fig. 4b). As a comparison, we applied Monocle3[28–33], a popular single-cell pipeline for cell lineage analysis. The resulting cell subpopulation network (Fig. 4c) is different from the actual lineage. Thus, SifiNet outperforms Monocle3 in SD2 to reveal cell developmental lineage.

**Figure 4:**
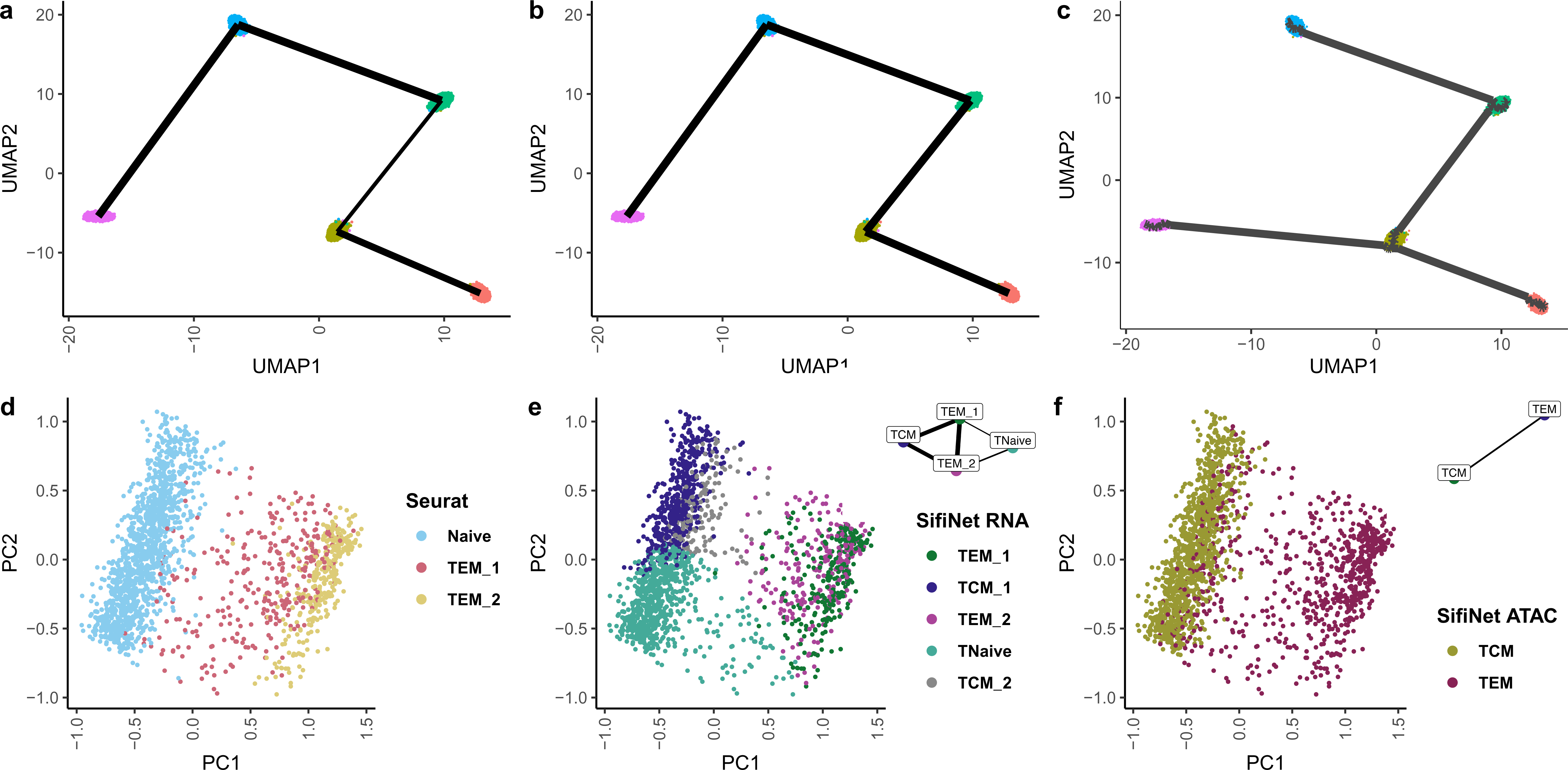
**(a)-(c) UMAPs of SD2**, with (a) the SifiNet-built lineage, (b) the true cell subpopulation lineage, and (c) the Monocle3-built lineage. **(d)-(f) Cellular GSVA score PC plots for CD8.** The x- and y-axis are PC directions of the GSVA scores based on the SifiNet-derived gene sets using scRNA-seq and scATAC-seq data. The cells were annotated by (d) Seurat on the multiome data (e) SifiNet on the scRNA-seq data (f) SifiNet on the scATAC-seq data. Cell sub-population networks are shown in (e) and (f).

### SifiNet accurately annotated CD8 T cells and identified long-lasting T cells

SifiNet is able to differentiate cells into subpopulations based on either single-cell transcriptomic or epigenomic data. Moreover, it can provide useful insights into the relationships between cell subpopulations.

In the CD8 scRNA-seq data, SifiNet identified four gene sets and annotated the cells with five cell subpopulations. Set 1 and Set 3 share *Gzma*, *Klrd1*, and *Klrg1*, markers for effector T cells[80,81]. Set 2 contains many central memory T cell (TCM) markers, including *Foxo1*, *Ccr7*, *Foxp1*, and *Tcf7* [82,83]. Set 4 contains 19 RPL and RPS genes, which are highly expressed in Näıve T cells[84]. Then, SifiNet clustered the cells into five subpopulations, TEM 1, TCM 1, TEM 2, TNäıve, and unassigned (the remaining cells), which are between the TCM 1 and TEM 2 cells (Fig. 4e). DEG analysis shows that the expression levels of TEM and TCM markers, *Gzma*, *Klrd1*, *Klrg1*, *Foxo1*, *Ccr7*, *Foxp1*, and *Tcf7*, in the unassigned subpopulation exhibit greater similarity to the TCM cells (Suppl. Fig. 10). Thus, we annotate the unassigned subpopulation as “TCM 2”. Central memory T cells (TCM 1, TCM 2) and Näıve T cells (TNaive) were previously annotated by Seurat as Näıve cells, and the TCM cells are missing based on Seurat annotations (Fig. 4d). This suggests that SifiNet has a stronger ability in differentiating cells. Further, SifiNet identified two effector T cell (TEM 1, TEM 2) subpopulations, both exhibiting bridging and transitional characteristics between TCM and TNaive cells (Fig. 4e). These results suggest CD8 subpopulation lineage relationships.

Previous studies used gene chromatin accessibility profiles to supplement genes’ expression profiles for defining T cell subpopulations.[85–87]. We adopted the same strategy. In the CD8 scATAC-seq data, SifiNet identified two feature gene sets. Set 1 includes TCM markers, such as *Klf7* and *Foxp1* [83]. Set 2 includes TEM markers, such as *Klrg1* [80]. These gene sets separated the cells into TCM and TEM cells, which shows excessive chromatin accessibility on TCM and TEM marker genes. Only one cell is in the unassigned subpopulation, which is not shown in the UMAP. In particular, we identified some CD8 cells with a TCM cell transcriptome profile but also excessive chromatin accessibility on TEM markers (Fig. 4e,f). They were considered as the most long-lasting T cells after yellow fever vaccination in a recent study[88]. Thus, further studying this CD8 cell subpopulation may lead to new insights into long-term immune protection.

### SifiNet scales to million-cell datasets

The rapid development of single-cell technology is leading to the generation of large-scale single-cell data collection, making more efficient computational methods necessary. Thus, while SifiNet primarily focuses on exploring subtle differences between similar cell subpopulations, we also applied SifiNet to a super-large-scale scRNA-seq dataset, MassiveRNA, to test its computational performance. MassiveRNA contains 10,448 genes and 1,287,071 cells after preprocessing. SifiNet took 12.2 hours and 101 GB peak memory to complete the analysis. To learn the impact of cell numbers on time cost and peak memory usage, we down-sampled MassiveRNA dataset into smaller datasets with cell numbers ranging between 10,000 to 1,000,000. SifiNet analysis on these datasets showed that the log10 running time is roughly linear with the log10 cell numbers (Suppl. Fig. 11). When the cell number is less than 50,000, the peak memory usage was approximately 4.1 GB, dominated by the multiple testing procedure. As the cell number increases above 50,000, the log10 peak memory usage was also roughly linear with the log10 cell number; the peak memory usage was reached in the co-expression calculation step (Suppl. Fig. 11). Overall, SifiNet is efficient with respect to computation time and memory usage. In general, the computation complexity of SifiNet is *O* (*Np*^2^), where *N* and *p* are the numbers of cells and genes after preprocessing.

### Discussion

Cell subpopulation annotation and understanding their relationships pose challenges, particularly when cells exhibit complex and subtle heterogeneity in states such as developmental stages, cell cycles, and senescence levels. While traditional clustering methods perform well in annotating subpopulations when there are clear functional distinctions reflected in cellular transcriptome profiles, they are less reliable in the presence of nuanced heterogeneity. To address these limitations, traditional methods often rely on domain-knowledge-assisted approaches. For instance, Seurat utilizes known cell cycle markers to derive cell cycle scores. However, for less understood cell statuses like senescence, the availability of marker genes may be limited. SifiNet demonstrates significant potential in these areas to identify new cell status markers and their associated cells.

Although SifiNet is computationally feasible for million-cell datasets, we do not recommend applying SifiNet directly on large-scale cell atlases, where the interested heterogeneity source is likely to be masked by other confounder heterogeneity sources. SifiNet’s strength lies in identifying the leading heterogeneity source in cell subpopulations with complex heterogeneity structures. Thus, for large-scale cell atlases, we suggest using UMAP, t-distributed Stochastic Neighbor Embedding (t-SNE)[89], or clustering methods to roughly cluster cells into visually obvious cell subpopulations where the interested heterogeneity is the leading. Then, SifiNet can be applied to the cell subpopulations to extract the feature gene sets of interest.

After SifiNet identifies the feature gene sets and the corresponding enriched cell subpopulation, we can immediately use the feature gene sets to annotate the cells. We can examine the identified feature genes and map them to the known marker genes in the public knowledge database. If we find a consensus among most feature genes, we will annotate the cell subpopulation with the corresponding known cell subpopulation. Some tools can help automate this process. For example, a recent study shows that GPT-4, a multi-modality large language model, can assist human researchers in accurately annotating known cell types based on a given set of feature genes[90]. If we cannot find a consensus in mapping the feature genes, the corresponding cell subpopulation is likely new. We can then use the feature genes and their co-expression network to annotate the cell type. We can also study the cell subpopulation network to understand the relationships between the cell subpopulations and other known cell subpopulations.

Moving forward, there are several avenues for further methodological improvement. Firstly, SifiNet’s performance is influenced by the quality of the gene co-expression network. If the interested cell population is very small, say, taking less than 5% of the total population, the current procedure to estimate the co-expression network might not be accurate enough for downstream analysis. Thus, we need new methods to enhance co-expression matrix estimation. Secondly, SifiNet utilizes the Louvain method with default resolution parameters to cluster feature genes, which generally works well when gene sets are well-separated. However, for gene sets with less distinct boundaries, optimizing gene set clustering becomes crucial to further enhance SifiNet’s performance on more challenging datasets. Thirdly, the incorporation of domain-knowledge-assisted approaches into SifiNet is being considered. This would allow SifiNet to leverage partially known marker genes, accurately capturing more feature genes and identifying corresponding cell subpopulations.

## Data availability

Monoclonal dataset is available at the Genome Sequencing Archive (GSA) with accession CRA008966. IPF1 and IPF2 datasets are collected from study GSE135893. Bone-Marrow dataset is collected from study GSE72857. CD8 dataset is available at 10x geneomics website https://support.10xgenomics.com/single-cell-multiome-atac-gex/datasets/1.0.0/pbmc_granulocyte_sorted_10k, and is also included in SeuratData R package as pbmc.rna and pbmc.atac. Seurat annotations for CD8 are available in SeuratData R package v0.2.2. MassiveRNA is downloaded from 10x genomics website https://support. 10xgenomics.com/single-cell-gene-expression/datasets/1.3.0/1M_neurons.

## Code availability

SifiNet v1.9 is used for the analysis in this study. SifiNet package is available at https://github.com/jichunxie/SifiNet. The codes used for generating the results shown in this paper are available at https://github.com/jichunxie/SifiNet_manu_support. GeneScape v1.0 used for generating numerical datasets is available on R CRAN.

## Acknowledgement

The authors thank the Duke CHSI for providing the computation resources.

