## Supplementary Figures and Supplementary Notes for "SifiNet: A robust and accurate method to identify feature gene sets and annotate cells"

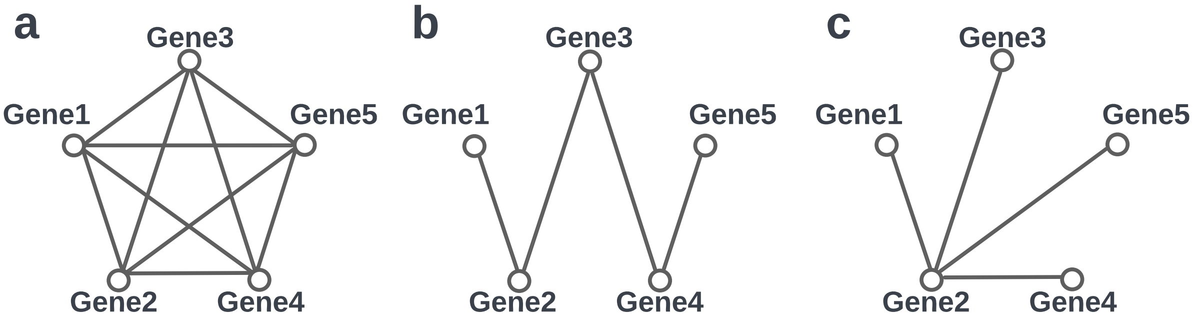


Supplementary Figure 1: **An illustration of co-expression network topology patterns for feature genes, pathway genes and hub genes.** **(a)** All feature genes tend to connect with each other, exhibiting strong community patterns. **(b)** Pathway genes usually co-express with several upstream or downstream genes. However, for far-away genes on the same pathway, the co-expressions quickly decay to almost zero. **(c)** Hub genes may regulate many genes, but these downstream genes do not exhibit obvious co-expression patterns.

a

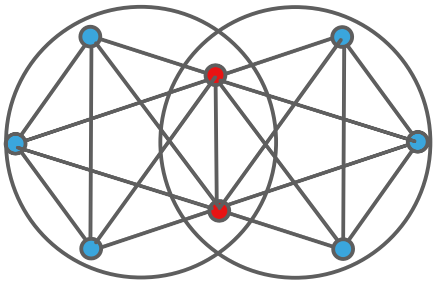

b

| | Core | Non-core |
|---|---|---|
| $2^{nd}$ | 1 | 0.571 |
| $3^{rd}$ | 0.679 | 0.679 |

Supplementary Figure 2: **An illustration of co-expression network topology patterns for core feature genes and non-core feature genes**. **(a)** Genes within the same circle belong to the same feature gene set. Core feature genes (blue) correspond to those genes unique to their corresponding feature gene sets. Non-core feature genes (red) are likely to be shared by different gene sets. **(b)** $2^{nd}$ and $3^{rd}$ connectivity values for core and non-core feature genes in (a).

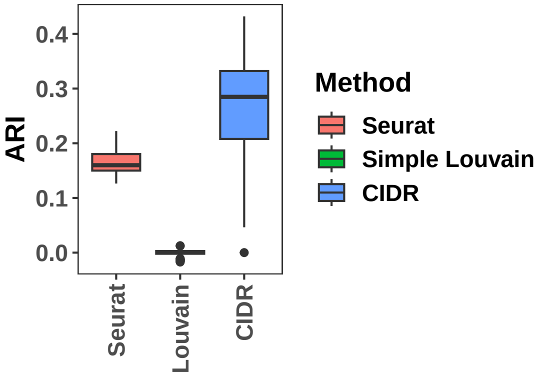


Supplementary Figure 3: **Cluster performance of SD1**. The performance of Seurat, simple Louvain and CIDR clustering at default resolution on SD1 evaluated by ARI[1].

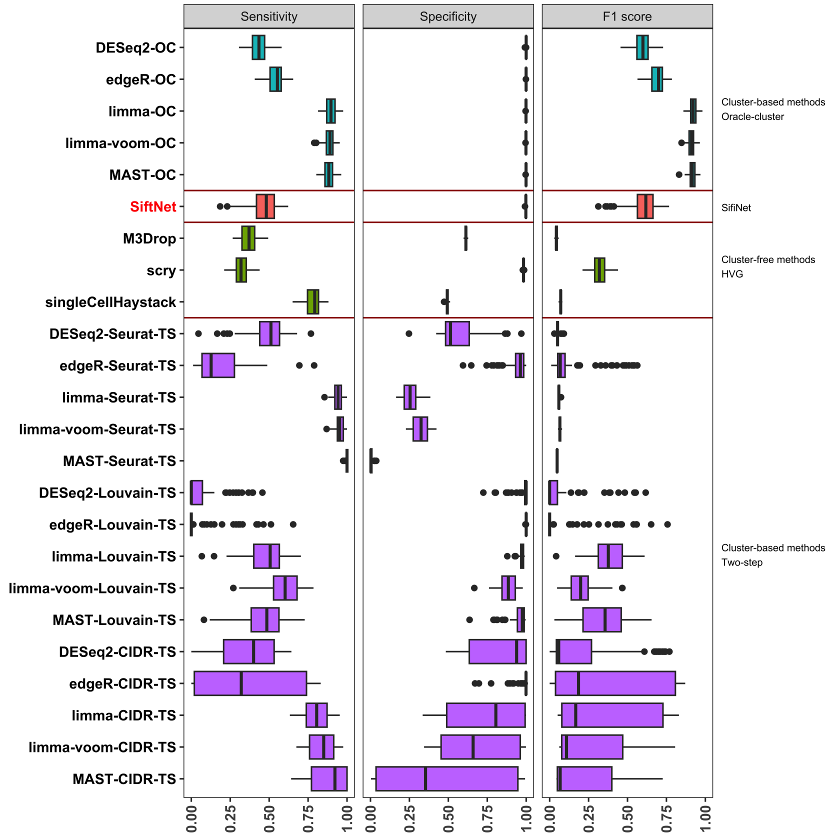


Supplementary Figure 4: **Feature gene detection performance in SD1**. The performance of feature gene detection methods: oracle-cluster methods ("OC" suffix, blue), SifiNet (red), HVG identification methods (green) and two-step methods ("TS" suffix, purple) on SD1 dataset. Methods shown above SifiNet are oracle methods for reference, while methods shown below SifiNet are realizable methods in practice.

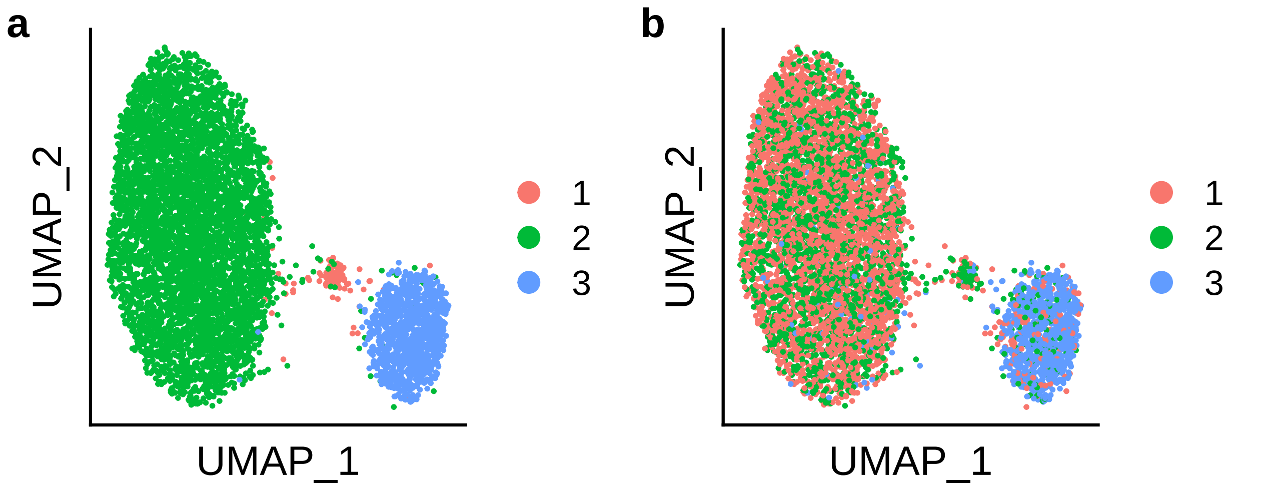


Supplementary Figure 5: **Cell clusters in SD1**. The UMAP plots of one SD1 replication colored by cell clusters from **(a)** the oracle-cluster approach (true cell type labels) and **(b)** CIDR with default resolution.

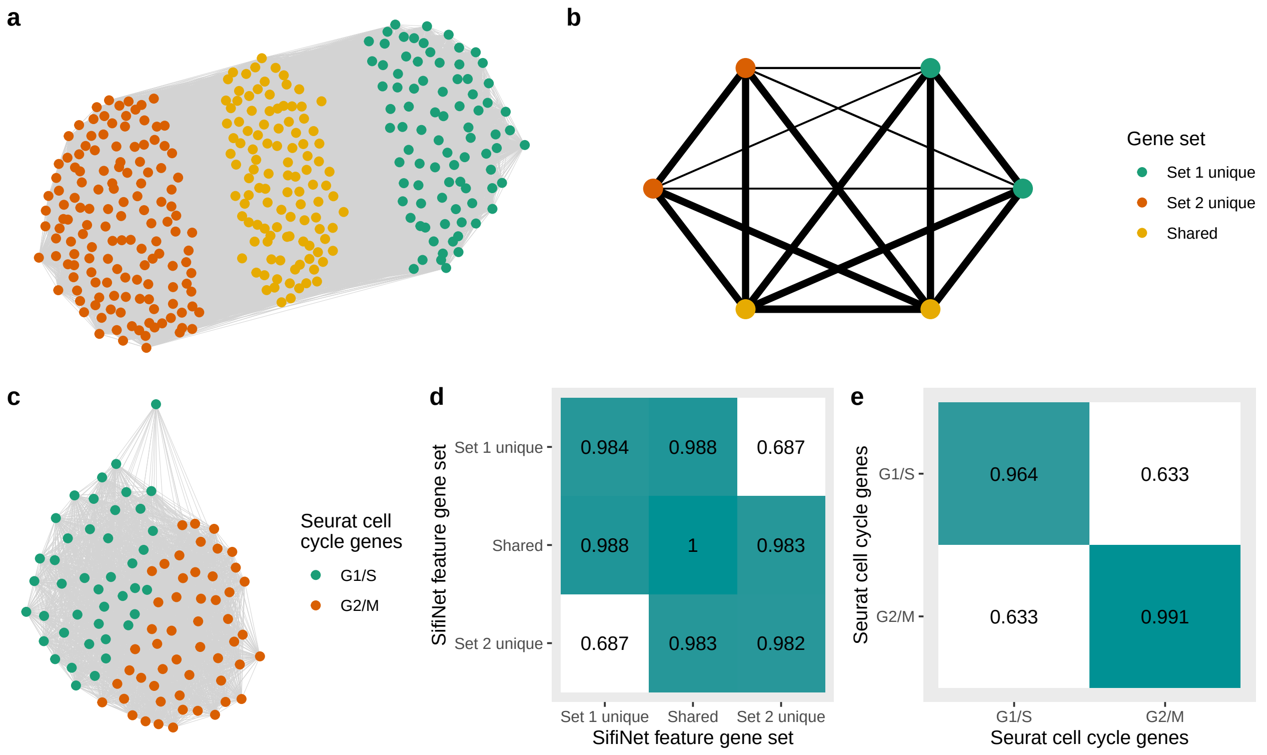


Supplementary Figure 6: **Gene co-expression network in Monoclonal.** **(a)** Co-expression between all feature genes identified by SifiNet. **(b)** Co-expression between feature genes unique to only one feature gene set (unique feature genes) and feature genes shared by the two feature gene sets. To better visualize the connections between feature gene sets, we randomly divided the Set 1 unique genes, Set 2 unique genes, and shared feature genes, each with two subsets. Then, we plotted the gene set network similar to SifiNet function 3. The edge widths between two gene sets represent their collective connectivity. Both the Set 1 unique and Set 2 unique gene subsets have a close connection with shared gene subsets. However, Set 1 unique and Set 2 unique gene subsets are less connected. **(c)** Co-expression between the known cell cycle markers used to calculate the Seurat cell cycle scores. **(d)** Collective connectivity (the number of edges between genes over the maximum possible number of edges) between feature gene sets identified by SifiNet. **(e)** Collective connectivity between the known cell cycle markers used to calculate the Seurat cell cycle scores.

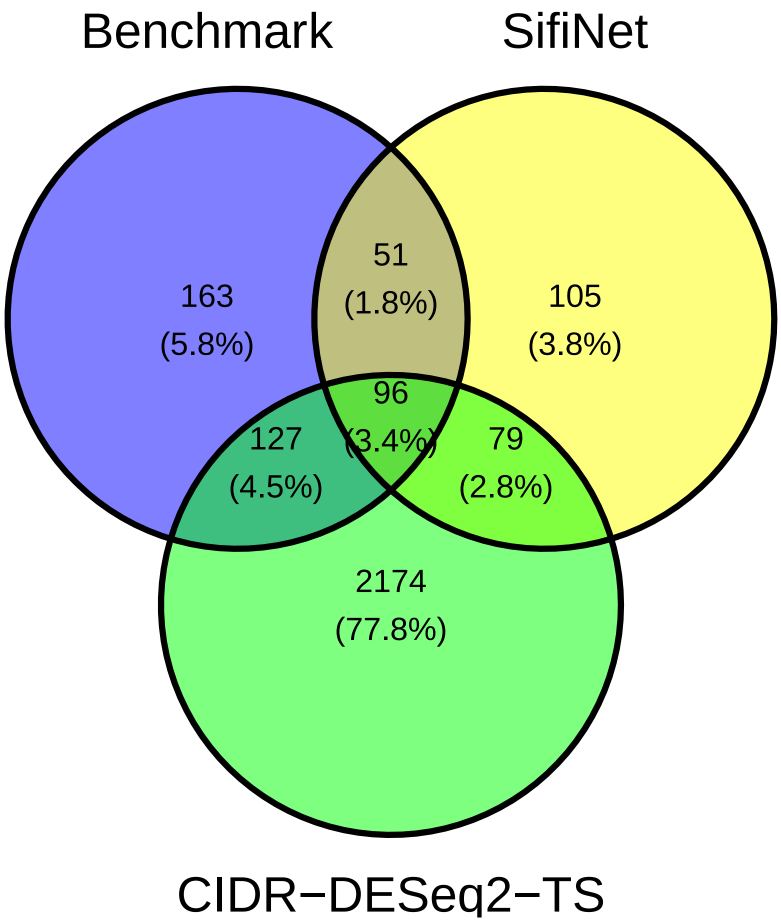


Supplementary Figure 7: **Venn diagram of benchmark genes, feature genes identified by SifiNet, and feature genes identified by CIDR-DESeq2-TS in Monoclonal.**

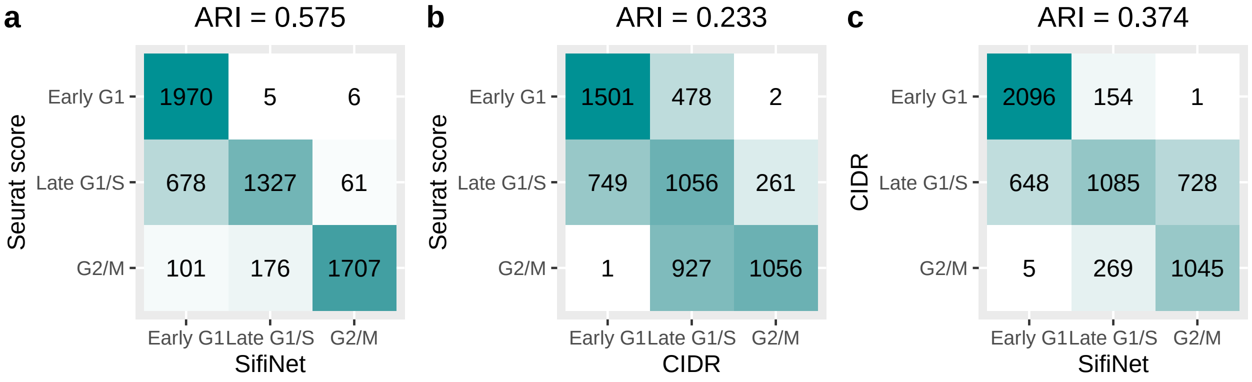


Supplementary Figure 8: **Confusion matrix to compare cell annotations for Monoclonal**. (a) SifiNet v.s. Seurat cell cycle score, (b) CIDR v.s. Seurat cell cycle score, (c) SifiNet v.s. CIDR. SifiNet has a higher ARI in cell annotations than CIDR.

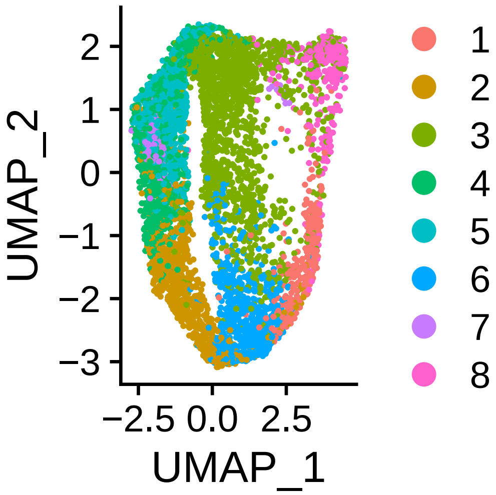


Supplementary Figure 9: **CIDR clustering at default resolution in Monoclonal**.

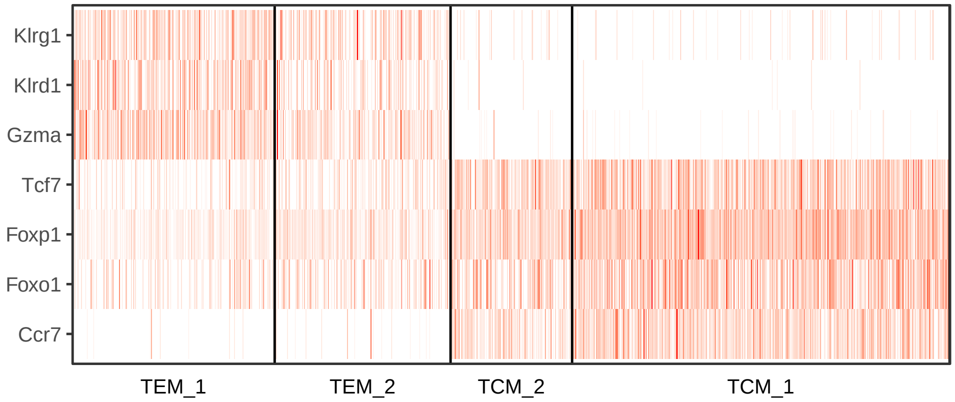


Supplementary Figure 10: **Heatmap of some marker gene expressions in the annotated cell subpopulations in CD8.** .

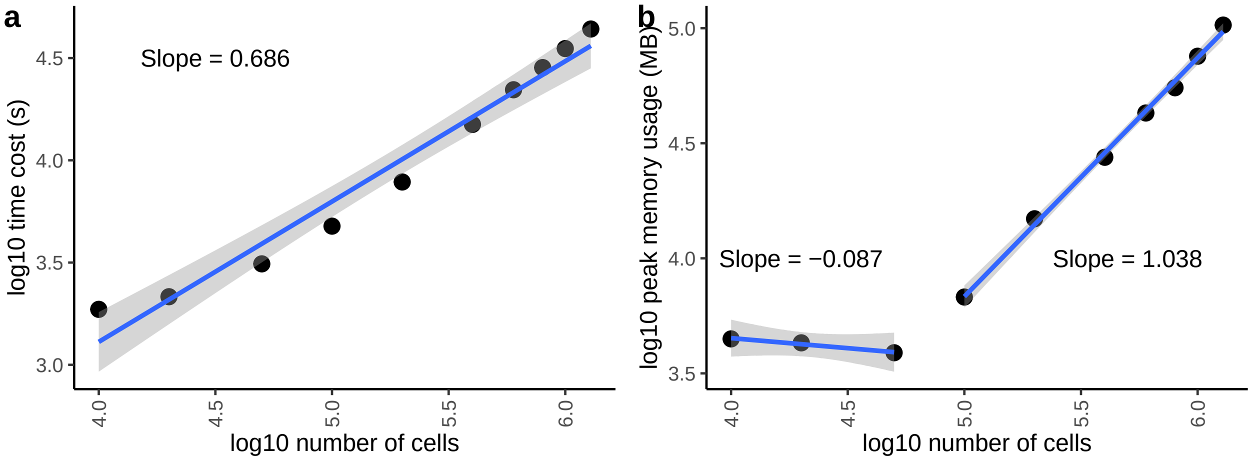


Supplementary Figure 11: **Time cost and peak memory usage on large dataset**. Time cost and peak memory usage of SifiNet on whole and downsampled MassiveRNA.

### Supplementary Note 1: Co-expression measures $D_{ij}$ and their normalized estimators $S_{ij}$ under various cases

#### Mathematical derivations of co-expression measures $D_{ij}$ under various cases

Feature genes are those over-expressed within a cell subpopulation. Its strict mathematical definition is given in Definition 1.

**Definition 1** (Feature gene)**.** Gene $i$ is called the feature gene of the cell subpopulation $\mathcal{C}$ if $P(Y_{ic} \leq y \mid c \in \mathcal{C}) \leq P(Y_{ic} \leq y \mid c \neq \mathcal{C})$ for any $y$, and $P(Y_{ic} \leq y_0 \mid c \in \mathcal{C}) < P(Y_{ic} \leq y_0 \mid c \neq \mathcal{C})$ for some $y_0$.

Recall that $D_{ij}$ is the co-expression measure between gene $i$ and gene $j$,

$$D_{ij} = \frac{1}{N} \sum_{c} P(Y_{ic} \leq q_{ic}, Y_{jc} \leq q_{jc} \mid s_c) - \tau_i \tau_j.$$

where $q_{ic}$ and $q_{jc}$ is $\tau_i$ and $\tau_j$ level cell-specific conditional quantiles of gene $i$ and gene $j$. Assume $Y_{ic} \perp\!\!\!\perp Y_{jc} \mid c \in \mathcal{C}_t, \ \forall, i, j, t$.

##### Case 1: Two feature genes over-expressing in the same cell subpopulation

Suppose gene $i$ and gene $j$ are both feature genes of $\tilde{\mathcal{C}}$, a subpopulation of cells. Let $q_{ic}$ and $q_{jc}$ be the cell-population-level $\tau_i$-th and $\tau_j$-th conditional quantile of gene $i$ and gene $j$ given $s_c$.

Based on Definition 1, feature genes over-express in the corresponding cell subpopulation. Therefore, when $c \in \tilde{\mathcal{C}}$, $P(Y_{ic} \leq q_{ic} | c \in \tilde{\mathcal{C}}) < \tau_i$, and $c \in \tilde{\mathcal{C}}$, $P(Y_{jc} \leq q_{jc} | c \in \tilde{\mathcal{C}}) < \tau_j$. Therefore, we have

$$\begin{aligned} P(Y_{ic} \leq q_{ic} | c \in \tilde{\mathcal{C}}) &= \tau_i - a_{ic}, \quad P(Y_{ic} \leq q_{ic} | c \notin \tilde{\mathcal{C}}) = \tau_i + a_{ic}, \\ P(Y_{jc} \leq q_{jc} | c \in \tilde{\mathcal{C}}) &= \tau_j - a_{jc}, \quad P(Y_{jc} \leq q_{jc} | c \notin \tilde{\mathcal{C}}) = \tau_j + a_{jc}, \end{aligned}$$

where $a_{ic} > 0$ is the differences between $P(Y_{ic} \leq q_{ic} | c \in \tilde{\mathcal{C}})$ and $\tau_i$, and $a_{jc} > 0$ is the differences between $P(Y_{jc} \leq q_{jc} | c \in \tilde{\mathcal{C}})$ and $\tau_j$.

Because

$$\begin{aligned} \tau_i &= \frac{1}{N} \sum_{c \in \tilde{\mathcal{C}}} P(Y_{ic} \leq q_{ic} | c \in \tilde{\mathcal{C}}) + \frac{1}{N} \sum_{c \notin \tilde{\mathcal{C}}} P(Y_{ic} \leq q_{ic} | c \notin \tilde{\mathcal{C}}) \\ &= \frac{1}{N} \sum_{c \in \tilde{\mathcal{C}}} (\tau_i - a_{ic}) + \frac{1}{N} \sum_{c \notin \tilde{\mathcal{C}}} (\tau_i + a_{ic}) = \tau_i - \sum_{c \in \tilde{\mathcal{C}}} a_{ic} + \sum_{c \notin \tilde{\mathcal{C}}} a_{ic}, \end{aligned}$$

we have $\sum_{c \in \tilde{\mathcal{C}}} a_{ic} = \sum_{c \notin \tilde{\mathcal{C}}} a_{ic}$. Similarly, $\sum_{c \in \tilde{\mathcal{C}}} a_{jc} = \sum_{c \notin \tilde{\mathcal{C}}} a_{jc}$. Then,

$$\begin{aligned} D_{ij} &= \frac{1}{N} \sum_{c \in \tilde{\mathcal{C}}} (\tau_i - a_{ic})(\tau_j - a_{jc}) + \frac{1}{N} \sum_{c \notin \tilde{\mathcal{C}}} (\tau_i + a_{ic})(\tau_j + a_{jc}) - \tau_i \tau_j \\ &= \frac{1}{N} \left( \sum_{c \in \tilde{\mathcal{C}}} a_{ic} a_{jc} + \sum_{c \notin \tilde{\mathcal{C}}} a_{ic} a_{jc} \right) > 0 \end{aligned} \tag{1}$$

Thus, these two genes have positive co-expression.

**Case 2: Two feature genes over-expressing in different cell subpopulations**

Suppose gene $i$ is feature gene of $\tilde{\mathcal{C}}_1$ and gene $j$ is feature gene of $\tilde{\mathcal{C}}_2$. Let

$$\begin{aligned} P(Y_{ic} \le q_{ic} | c \in \tilde{\mathcal{C}}_1) = \tau_i - a_{ic}, \quad P(Y_{ic} \le q_{ic} | c \notin \tilde{\mathcal{C}}_1) = \tau_i + a_{ic}, \\ P(Y_{jc} \le q_{jc} | c \in \tilde{\mathcal{C}}_2) = \tau_j - a_{jc}, \quad P(Y_{jc} \le q_{jc} | c \notin \tilde{\mathcal{C}}_2) = \tau_j + a_{jc}, \end{aligned}$$

where $a_{ic} > 0$ and $a_{jc} > 0$. We have

$$\sum_{c \in \tilde{\mathcal{C}}_1} a_{ic} = \sum_{c \notin \tilde{\mathcal{C}}_1} a_{ic}, \quad \sum_{c \in \tilde{\mathcal{C}}_2} a_{jc} = \sum_{c \notin \tilde{\mathcal{C}}_2} a_{jc} \tag{2}$$

Then,

$$\begin{aligned} D_{ij} =& \frac{1}{N} \sum_{c \in \tilde{\mathcal{C}}_1 \backslash \tilde{\mathcal{C}}_2} (\tau_i - a_{ic})(\tau_j + a_{jc}) + \frac{1}{N} \sum_{c \in \tilde{\mathcal{C}}_2 \backslash \tilde{\mathcal{C}}_1} (\tau_i + a_{ic})(\tau_j - a_{jc}) \\ &+ \frac{1}{N} \sum_{c \in \tilde{\mathcal{C}}_2 \cap \tilde{\mathcal{C}}_1} (\tau_i - a_{ic})(\tau_j - a_{jc}) + \frac{1}{N} \sum_{c \in (\tilde{\mathcal{C}}_1 \cup \tilde{\mathcal{C}}_2)^c} (\tau_i + a_{ic})(\tau_j + a_{jc}) - \tau_i \tau_j \\ =& \frac{1}{N} \left\{ - \sum_{c \in \tilde{\mathcal{C}}_1 \backslash \tilde{\mathcal{C}}_2} a_{ic} a_{jc} - \sum_{c \in \tilde{\mathcal{C}}_2 \backslash \tilde{\mathcal{C}}_1} a_{ic} a_{jc} + \sum_{c \in \tilde{\mathcal{C}}_1 \cap \tilde{\mathcal{C}}_2} a_{ic} a_{jc} + \sum_{c \in (\tilde{\mathcal{C}}_1 \cup \tilde{\mathcal{C}}_2)^c} a_{ic} a_{jc} \right\}. \end{aligned}$$

Under general cases, when two genes over-express in different cell populations, they may have either positive or negative co-expression.

However, when $\tilde{\mathcal{C}}_1 \cap \tilde{\mathcal{C}}_2 = \emptyset$,

$$D_{ij} = \frac{1}{N} \left\{ - \sum_{c \in \tilde{\mathcal{C}}_1 \cup \tilde{\mathcal{C}}_2} a_{ic} a_{jc} + \sum_{c \in (\tilde{\mathcal{C}}_1 \cup \tilde{\mathcal{C}}_2)^c} a_{ic} a_{jc} \right\}. \tag{3}$$

If $\tilde{\mathcal{C}}_1 \cup \tilde{\mathcal{C}}_2$ is the entire cell population, *i.e.*, $(\tilde{\mathcal{C}}_1 \cup \tilde{\mathcal{C}}_2)^c = \emptyset$, then the second term is zero, and thus $D_{ij} < 0$. These two genes have negative co-expression.

Alternatively, when cell library sizes are similar, then all the $a_{ic}, a_{jc}$ are similar within each cell subpopulation. Further, by (2), we assume

$$a_{ic} \approx \frac{A_i}{|\tilde{\mathcal{C}}_1|} \text{ for } c \in \tilde{\mathcal{C}}_1, \quad a_{ic'} \approx \frac{A_i}{N - |\tilde{\mathcal{C}}_1|} \text{ for } c' \notin \tilde{\mathcal{C}}_1, \quad a_{jc} \approx \frac{A_j}{|\tilde{\mathcal{C}}_2|} \text{ for } c \in \tilde{\mathcal{C}}_2, \quad a_{ic'} \approx \frac{A_j}{N - |\tilde{\mathcal{C}}_2|} \text{ for } c' \notin \tilde{\mathcal{C}}_2.$$

Based on (3), we have

$$D_{ij} \approx \left\{ - \frac{1}{N - |\tilde{\mathcal{C}}_2|} - \frac{1}{N - |\tilde{\mathcal{C}}_1|} + \frac{N - |\tilde{\mathcal{C}}_1| - |\tilde{\mathcal{C}}_2|}{(N - |\tilde{\mathcal{C}}_1|)(N - |\tilde{\mathcal{C}}_2|)} \right\} A_i A_j < \left\{ - \frac{1}{N - |\tilde{\mathcal{C}}_1|} \right\} A_i A_j < 0.$$

Thus, these two genes have negative co-expression.

**Case 3: At least one gene is not a feature gene**

Without loss of generality, suppose gene $i$ is not a feature gene. Then in (1), we can set $a_{ic} = 0$ for all $c$. Then, $D_{ij} = 0$, *i.e.*, these two genes are independent.

#### Validating the co-expressions under various situations

In practice, the gene co-expression $D_{ij}$ is unknown. However, we can derive its standardized estimator of $S_{ij}$, as defined in Materials and Methods. We verified the mathematical derivation of $D_{ij}$ by checking the distribution of $S_{ij}$ of different gene pairs in an experimental dataset, TvB.

The dataset, TvB, is a subset of a single-cell multiome (gene expression and ATAC-seq) dataset from 10x Genomics[2]. We only used the gene expression part of this dataset. An annotation of cell types was provided by Seurat v4, available at `SeuratData` R package v0.2.2. To simplify the analysis, we only include T cells and B cells in the analysis. The T cell cohort includes "CD8 Naive" and "CD4 Naive" cells; and B cell cohort includes "Intermediate B", "Memory B", and "Naive B" cells. Those cells with total read count greater than 1.8 times of the mean total read count across all cells were removed. We further removed cells with fewer than 2% of the genes detected, and then the genes detected in fewer than 2% of the cells. No additional normalization was performed before we applied SifiNet.

We curated marker gene sets for T cells and B cells from the GSEA MSigDB[3–5], respectively. We downloaded the human C7 immunologic signature gene sets v2023.1 and extract the gene sets with keywords "TCELL_VS_BCELL_UP" or "TCELL_VS_BCELL_DN" in their names[6,7]. Genes that were included in at least 2 studies were selected. In total, forty-six "TCELL_VS_BCELL_UP" genes were included in the T-cell marker gene set, and seventy "TCELL_VS_BCELL_DN" genes were included in the B-cell marker gene set.

Next, we calculated $S_{ij}$ for all the gene pairs, and devided them into four groups: (1) $S_{ij}$ between two genes in the same marker gene set; (2) $S_{ij}$ between two genes in different marker gene sets; (3) $S_{ij}$ between one gene in a marker gene set and one gene not in any marker gene set; (4) $S_{ij}$ between two genes not in any marker gene set. The distributions of $S_{ij}$ in these four groups are shown in Suppl. Fig. 1. The results verified our mathematical derivations. For genes in the same marker gene set, $S_{ij}$ have positive expectation; for two genes in different marker gene sets, $S_{ij}$ have negative expectation; if at least one of the genes is not a feature gene, $S_{ij}$'s expectation is close to 0, and the distribution of $S_{ij}$ is close to the reference standard Gaussian distribution.

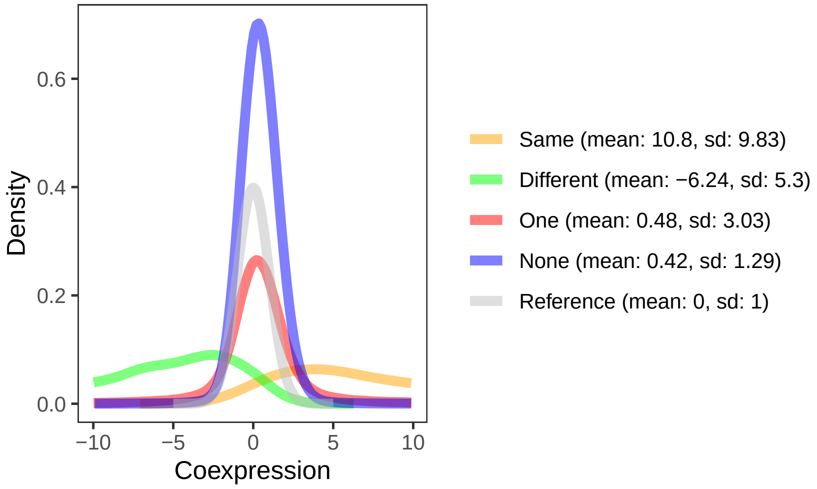


Note Figure 1: **Density of quantile association estimator $S_{ij}$ in TvB**. Same (orange): density of $S_{ij}$ when two feature genes are over-expressed in the same cell subpopulation (both T cells or both B cells); Different (green): density of $S_{ij}$ when two feature genes are over-expressed in the different cell subpopulation (one in T cells and one in B cells); One (red): density of $S_{ij}$ when one of the two genes is a feature gene while the other is not a feature gene; None (blue): density of $S_{ij}$ when neither of the two genes are feature genes; Reference (gray): The density of standard Gaussian distribution.

### Supplementary Note 2: Asymptotic null distribution of $S_{ij}$ when at least one gene is not a feature gene.

This note aims to derive the asymptotic distribution of $S_{ij}$ when $i$ or $j$ is not a feature gene, where $S_{ij}$ is defined in (1) in Method. Here, "asymptotics" means that the number of cell $N$ goes to infinity.

**Theorem 1.** *Recall that* $\mathrm{H}_{null,ij}: \; Y_{ic} \perp\!\!\!\perp Y_{jc} \mid s_c$*, for all* $i, j \in \mathcal{V}$ *and* $i < j$*. Let* $\mathbb{D}_0 = \{(i,j) : 1 \le i < j \le p, \mathrm{H}_{null,ij}$ *is true*$\}$*. Under conditions 1-4, for any* $C_0$*, when* $0 \le t \le C_0 \log(n \vee p)^{1/2}$*, we have*

$$\lim_{N\to\infty} \sup_{(i,j)\in\mathbb{D}_0} \left| \frac{P(|S_{ij}| \ge t)}{2\bar{\Phi}(t)} - 1 \right| = 0.$$

Theorem 1 builds the theoretical foundation of the paper. It shows that under the null, $|S_{ij}|$ asymptotically converges to $|Z|$, where $Z \sim N(0,1)$. This result is useful in performing multiple testing on the edges of the co-expression graph. The proof of Theorem 1 requires 4 mild conditions, which are introduced in the following "Theorem, lemmas, and proofs" section.

We designed a T cell versus B cell experiment to check the gene co-expression estimator distributions under different situations (Suppl. Note 1). When none or only one gene is the feature gene, the distribution of $S_{ij}$ is close to the standard Gaussian distribution (Note Fig. 1). Additionally, a simple simulation study to show the asymptotic null distribution is shown in Note Fig. 2.

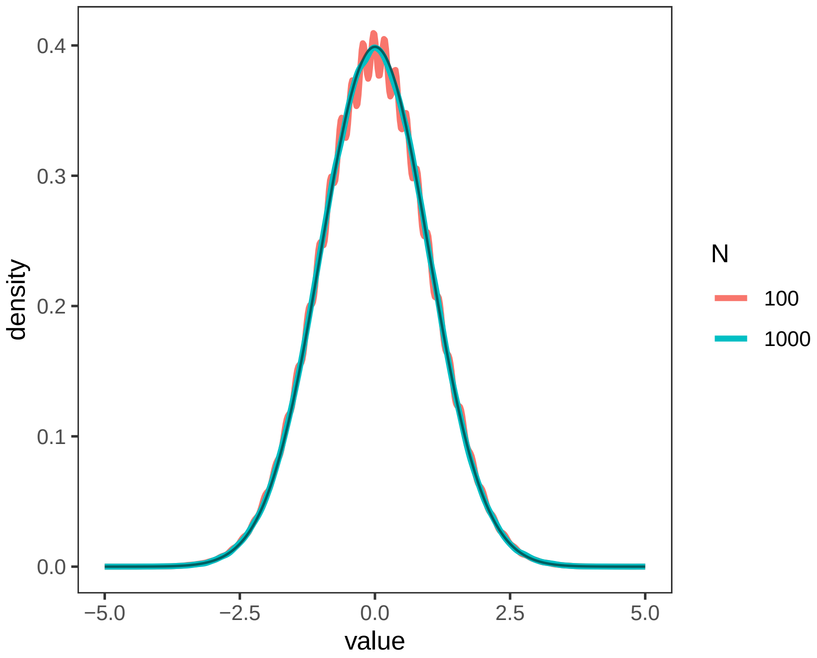


Note Figure 2: **Asymptotic null distribution of $S_{ij}$ on simulated dataset**. The red and green curves are the null density of $S_{ij}$ when $N = 100$ and $N = 1000$, respectively; while the black curve is the density of standard Gaussian distribution. When $N = 1000$, the null distribution of $S_{ij}$ is very close to the standard Gaussian density.

The complementary cumulative density function of the standard Gaussian, denoted by, $\bar{\Phi}(t)$, also converges to 0 as $t$ goes to infinity. To ensure the asymptotic false discovery rate control in the following multiple testing procedure, Theorem 1 also specifies that the CDF difference over $2\bar{\Phi}(t)$ also converges to zero, as $t$ in a range slowly diverge with $N$ as $N$ goes to infinity.

#### Theorem, lemmas, and proofs

Assume all cells are all independent. For the same cell, we assume the gene feature values are independent conditioning on cell subpopulations,

$$Y_{ic} \perp\!\!\!\perp Y_{jc} \mid c \in \mathcal{C}_t, \ \forall, i, j, t.$$

Recall that $D_{ij}$'s standardized estimator $S_{ij}$ is defined as

$$S_{ij} = \frac{\sum_{c=1}^{N}[I(Y_{ic} \le \hat{q}_{ic}, Y_{jc} \le \hat{q}_{jc}) - \hat{\tau}_i\hat{\tau}_j]}{\sqrt{N\hat{\tau}_i(1-\hat{\tau}_i)\hat{\tau}_j(1-\hat{\tau}_j)}}, \tag{4}$$

where $\hat{q}_{ic}$ is the estimated (by quantile regression) cell-specific quantile of gene $i$ at the percentile $\tau_i$, and $\hat{\tau}_i = \sum_{c=1}^{N} I(Y_{ic} \le \hat{q}_{ic})/N$ is the realized percentile. To show the asymptotic properties of $S_{ij}$, we define two intermediate quantities:

$$A_{ij} = \frac{\sum_{c=1}^{N}[I(Y_{ic} \le q_{ic}) - \tau_i][I(Y_{jc} \le q_{jc}) - \tau_j]}{\sqrt{N\tau_i(1-\tau_i)\tau_j(1-\tau_j)}}, \ B_{ij} = \frac{\sum_{c=1}^{N}[I(Y_{ic} \le \hat{q}_{ic}, Y_{jc} \le \hat{q}_{jc}) - \hat{\tau}_i\hat{\tau}_j]}{\sqrt{N\tau_i(1-\tau_i)\tau_j(1-\tau_j)}}.$$

Here, $q_{ic} = q_i(s_c)$ is the true cell-specific $\tau_i$-quantile of gene $i$.

To derive the asymptotic properties introduced in Theorem 1, we need the following conditions:

1. Assume the conditional probability density function $f_{\mathbf{s},i}$ of $Y_{ic}|s_c$ is bounded, i.e., $|f_{\mathbf{s},i}(y)| \le C_1$, for $\forall y$, $s_c \in \mathbb{R}$, $i \in [p]$.

2. Assume $\exists C$ and some small constant $\epsilon > 0$, such that

$$P\left\{|\hat{q}_{ic} - q_{ic}| > 2C(1+\epsilon)N^{-1/2}log(N \vee p)^{1/2}\right\} < (N \vee p)^{-2-\epsilon}.$$

3. Assume $p \le N^r$ for some $r > 0$.

4. Assume $\tau_i$ to be bounded away from 0 and 1, i.e., $0 < \tau_{\min} \le \tau_i \le \tau_{\max} < 1$ for $\forall i \in [p]$.

Conditions 1 and 2 are mild conditions as shown in Xie and Li[8]. Condition 3 is generally satisfied in most cases. For condition 4, by default, we set $\tau_i = (1 + \hat{p}_{i0})/2$, where $\hat{p}_{i0} = \sum_{c=1}^{N} I(Y_{ic} = 0)/N$. In practice, condition 4 is satisfied by removing genes expressed in fewer than some proportion of the cells in the preprocessing procedure.

**Lemma 1.** *Let* $\Delta_{ij} = B_{ij} - A_{ij}$. *Then* $\sup_{(i,j)\in\mathbb{D}_0} E(|\Delta_{ij}|^m) \le C_m\{\log(N \vee p)^2/N\}^{m/2}$, *for some constant* $C_m$.

*Proof.* We employed the similar techniques when proving lemma 1 in Xie and Li[8], and devided $\Delta_{ij}$

into four parts $\Delta_{ij} = \sum_{l=1}^{4} \Delta_{l,ij}$, where

$$\Delta_{1,ij} = \frac{\sum_{c=1}^{N}[I(Y_{ic} \leq q_{ic}) - \tau_i][I(Y_{jc} \leq \hat{q}_{jc}) - I(Y_{jc} \leq q_{jc})]}{\sqrt{N\tau_i(1-\tau_i)\tau_j(1-\tau_j)}},$$

$$\Delta_{2,ij} = \frac{\sum_{c=1}^{N}[I(Y_{ic} \leq \hat{q}_{ic}) - I(Y_{ic} \leq q_{ic})][I(Y_{jc} \leq q_{jc}) - \tau_j]}{\sqrt{N\tau_i(1-\tau_i)\tau_j(1-\tau_j)}},$$

$$\Delta_{3,ij} = \frac{\sum_{c=1}^{N}[I(Y_{ic} \leq \hat{q}_{ic}) - I(Y_{ic} \leq q_{ic})][I(Y_{jc} \leq \hat{q}_{jc}) - I(Y_{jc} \leq q_{jc})]}{\sqrt{N\tau_i(1-\tau_i)\tau_j(1-\tau_j)}},$$

$$\Delta_{4,ij} = \frac{\sum_{c=1}^{N}\tau_i[I(Y_{jc} \leq \hat{q}_{jc}) - \tau_j] + \tau_j[I(Y_{ic} \leq \hat{q}_{ic}) - \tau_i] - [\hat{\tau}_i\hat{\tau}_j - \tau_i\tau_j]}{\sqrt{N\tau_i(1-\tau_i)\tau_j(1-\tau_j)}}.$$

By Theorem 2.2 in Koenker[9], suppose there are $N_0$ parameters included in the quantile regression. Then

$$\tau_i - \frac{N_0}{N} \leq \hat{\tau}_i \leq \tau_i + \frac{N_0}{N}.$$

Thus, we have

$$\Delta_{4,ij} = O\left(N^{-1/2}\right).$$

The rest of the proof is the same as lemma 1 in Xie and Li[8]. □

**Lemma 2.** *Let $\bar{\Phi}$ be the complementary CDF of standard Gaussian distribution. Then*

$$\lim_{t\to\infty} \frac{\bar{\Phi}(t)}{(t\sqrt{2\pi})^{-1}\exp(-t^2/2)} = 1$$

*Proof.* Suppose that $X$ follows a standard normal distribution $N(0,1)$. $\bar{\Phi} = P(X \geq t)$. As shown in Richter and Schumacher[10], based on the representation formula for Laplace integral.

$P(X \geq t) = \frac{1}{\sqrt{2\pi}}\int_t^\infty e^{-x^2/2}dx = \frac{t}{\sqrt{2\pi}}\int_1^\infty e^{-t^2x^2/2}dx$. Let $f(x) = 1$, $S(x) = -x^2/2$, $\lambda = t^2$, we have $\lim_{t\to\infty} \frac{P(X \geq t)}{(t\sqrt{2\pi})^{-1}\exp(-t^2/2)} = 1$. □

**Lemma 3.** *When $0 \leq t \leq C_0 \log(N \vee p)^{1/2}$, $\forall\epsilon > 0$, $\frac{P[|\Delta_{ij}| > \log(n \vee p)^{-1}]}{\bar{\Phi}(t)} \leq \epsilon$ for large enough $N$.*

*Proof.* By Markov's inequality and Lemma 1, $P(|\Delta_{ij}| > \log(N \vee p)^{-1}) \leq C_m \log(N \vee p)^{2m} N^{-m/2}$.

By assumption, $p \leq cN^r$. We have $N \vee p \leq (1+c)N^{1+r}$. By Lemma 2,

$\frac{P(|\Delta_{ij}| > \log(N \vee p)^{-1})}{\bar{\Phi}(t)} \leq \frac{C_m \log(N \vee p)^{2m} N^{-m/2}}{(C_0 \log(N \vee p)^{1/2}\sqrt{2\pi})^{-1}\exp(-C_0^2 \log(N \vee p)/2)}$

$\leq C'_m \log(N \vee p)^{2m+1/2} N^{-m/2+C_0^2(1+r)/2}$.

We may choose $m > C_0^2(1+r)$ such that $\frac{P(|\Delta_{ij}| > \log(N \vee p)^{-1})}{\bar{\Phi}(t)} \leq \epsilon$. □

**Lemma 4.** $\sup_{(i,j)\in\mathbb{D}_0} |\frac{P(|A_{ij}| \geq t)}{2\bar{\Phi}(t)} - 1| \leq C_2(1+t)^{3/2}N^{-1/2}$, *when $t = o(N^{1/3})$.*

*Proof.* By central limit theorem, $A_{ij}$ asymptotically follows the standard Gaussian distribution. The following proof is similar to Lemma 4 in Xie and Li[8]. □

*Proof of Theorem 1.* By the proof of Lemma 1 and condition 4,

$$\frac{\tau_i(1-\tau_i)\tau_j(1-\tau_j) - \hat{\tau}_i(1-\hat{\tau}_i)\hat{\tau}_j(1-\hat{\tau}_j)}{\hat{\tau}_i(1-\hat{\tau}_i)\hat{\tau}_j(1-\hat{\tau}_j)} = O\left(\frac{1}{N}\right).$$

Using Taylor expansion of $\sqrt{1+x}$, for sufficiently large $N$, there exists constant $C_3$ such that

$$\begin{aligned}
&|S_{ij} - B_{ij}| \\
=&|B_{ij}|\left|\sqrt{\frac{\tau_i(1-\tau_i)\tau_j(1-\tau_j)}{\hat{\tau}_i(1-\hat{\tau}_i)\hat{\tau}_j(1-\hat{\tau}_j)}} - 1\right| \\
=&|B_{ij}|\left|\frac{\tau_i(1-\tau_i)\tau_j(1-\tau_j) - \hat{\tau}_i(1-\hat{\tau}_i)\hat{\tau}_j(1-\hat{\tau}_j)}{2\hat{\tau}_i(1-\hat{\tau}_i)\hat{\tau}_j(1-\hat{\tau}_j)} + o\left(\frac{\tau_i(1-\tau_i)\tau_j(1-\tau_j) - \hat{\tau}_i(1-\hat{\tau}_i)\hat{\tau}_j(1-\hat{\tau}_j)}{\hat{\tau}_i(1-\hat{\tau}_i)\hat{\tau}_j(1-\hat{\tau}_j)}\right)\right| \\
\leq&\frac{C_3}{N}|B_{ij}|.
\end{aligned}$$

Then,

$$\begin{aligned}
&\sup_{(i,j)\in\mathbb{D}_0}\frac{P(|S_{ij}|>t)}{2\bar{\Phi}(t)} = \sup_{(i,j)\in\mathbb{D}_0}\frac{P(|A_{ij} + (B_{ij}-A_{ij}) + (S_{ij}-B_{ij})|>t)}{2\bar{\Phi}(t)} \\
\leq&\sup_{(i,j)\in\mathbb{D}_0}\frac{P(|A_{ij}|>t-2\log(N\vee p)^{-1}) + P(|\Delta_{ij}|>\log(N\vee p)^{-1}) + P(|S_{ij}-B_{ij}|>\log(N\vee p)^{-1})}{2\bar{\Phi}(t)} \\
\leq&\sup_{(i,j)\in\mathbb{D}_0}\frac{P(|A_{ij}|>t-2\log(N\vee p)^{-1}) + P(|\Delta_{ij}|>\log(N\vee p)^{-1}) + P(|B_{ij}|>\frac{N}{C_3}\log(N\vee p)^{-1})}{2\bar{\Phi}(t)} \\
\leq&\sup_{(i,j)\in\mathbb{D}_0}\frac{P(|A_{ij}|>t-2\log(N\vee p)^{-1}) + 2P(|\Delta_{ij}|>\log(N\vee p)^{-1}) + P(|A_{ij}|>(\frac{N}{C_3}-1)\log(N\vee p)^{-1})}{2\bar{\Phi}(t)} \\
=&\sup_{(i,j)\in\mathbb{D}_0}\frac{P(|A_{ij}|>t-2\log(N\vee p)^{-1})}{2\bar{\Phi}(t-2\log(N\vee p)^{-1})}\frac{2\bar{\Phi}(t-\log(N\vee p)^{-1})}{2\bar{\Phi}(t)} + \sup_{(i,j)\in\mathbb{D}_0}\frac{2P(|\Delta_{ij}|>\log(N\vee p)^{-1})}{2\bar{\Phi}(t)} \\
+&\sup_{(i,j)\in\mathbb{D}_0}\frac{P(|A_{ij}|>(\frac{N}{C_3}-1)\log(N\vee p)^{-1})}{2\bar{\Phi}((\frac{N}{C_3}-1)\log(N\vee p)^{-1})}\frac{2\bar{\Phi}((\frac{N}{C_3}-1)\log(N\vee p)^{-1})}{2\bar{\Phi}(t)}
\end{aligned}$$

Based on the continuity of $\bar{\Phi}$, for any small $\epsilon_p$, when $p$ is sufficiently large,

$$\left|\frac{\bar{\Phi}(t-\log(N\vee p)^{-1})}{\bar{\Phi}(t)} - 1\right| \leq \epsilon_p.$$

By Lemma 3,

$$\frac{2P(|\Delta_{ij}|>\log(N\vee p)^{-1})}{2\bar{\Phi}(t)} \leq 2\epsilon_p.$$

By Lemma 4,

$$\left|\frac{P(|A_{ij}|>(\frac{N}{C_3}-1)\log(N\vee p)^{-1})}{2\bar{\Phi}((\frac{N}{C_3}-1)\log(N\vee p)^{-1})} - 1\right| \leq \epsilon_p.$$

$$\left|\frac{P(|A_{ij}|>t-\log(N\vee p)^{-1})}{2\bar{\Phi}(t-\log(N\vee p)^{-1})} - 1\right| \leq \epsilon_p.$$

By Lemma 2,

$$\begin{aligned}
&\lim_{N\to\infty} \frac{2\bar{\Phi}((\frac{N}{C_3}-1)\log(N\vee p)^{-1})}{2\bar{\Phi}(t)} \\
\leq &\lim_{N\to\infty} \frac{2\bar{\Phi}((\frac{N}{C_3}-1)\log(N\vee p)^{-1})}{2\bar{\Phi}(C_0\log(n\vee p)^{1/2})} \\
= &\lim_{N\to\infty} \frac{C_0\log(n\vee p)^{3/2}}{(\frac{N}{C_3}-1)}\exp\{-\frac{[(\frac{N}{C_3}-1)^2 - C_0^2\log(n\vee p)^3]}{2\log(N\vee p)^2}\} \\
=&0.
\end{aligned}$$

Then,

$$\frac{2\bar{\Phi}((\frac{N}{C_3}-1)\log(N\vee p)^{-1})}{2\bar{\Phi}(t)} \leq \epsilon_p.$$

So,

$$\sup_{(i,j)\in\mathbb{D}_0} \frac{P(|S_{ij}|>t)}{2\bar{\Phi}(t)} \leq 1+5\epsilon_p.$$

Similarly, by

$$\begin{aligned}
&\frac{P(|S_{ij}|>t)}{2\bar{\Phi}(t)} = \frac{P(|A_{ij}+(B_{ij}-A_{ij})+(S_{ij}-B_{ij})|>t)}{2\bar{\Phi}(t)} \\
\geq &\frac{P(|A_{ij}|>t+2\log(N\vee p)^{-1}) - P(|\Delta_{ij}|>\log(N\vee p)^{-1}) - P(|S_{ij}-B_{ij}|>\log(N\vee p)^{-1})}{2\bar{\Phi}(t)}.
\end{aligned}$$

we can show that

$$\sup_{(i,j)\in\mathbb{D}_0} \frac{P(|S_{ij}|>t)}{2\bar{\Phi}(t)} \geq 1-5\epsilon_p.$$

□

### Supplementary Note 3: Some details of SifiNet algorithm

#### Sampling algorithm to calculate the 3rd-order connectivities

We randomly sample $m$ genes from gene $i$'s 2nd-order neighbors, with the sampling probabilities proportional to the number of the gene's connecting edges with the genes in $\mathcal{N}_i$. In other word, the weight of gene $k$ for calculating gene $i$'s 3rd-order connectivity $w_{ik} \propto |\{j : (k, j) \in \mathcal{E}, j \in \mathcal{N}_i\}|$. The resulting gene set at repetition $\ell$ is denoted by $\mathcal{A}_i^{(\ell)}$. Then, $C_{3i}$ computes the connectivity levels between the genes in $\mathcal{A}_i^{(\ell)}$ as $C_{3i} = \sum_{\ell=1}^{L} |\mathcal{E} \cap \{(j, k) : j, k \in \mathcal{A}_i^{(\ell)}\}|/Lm(m-1)$.

#### Multiple testing procedure to elucidate the co-expression graph

The gene co-expression graph by $\mathcal{G} = \{\mathcal{V}, \mathcal{E}\}$ with $\mathcal{V} = [p]$, and $\mathcal{E}$ denoting the edge set. To obtain $\mathcal{E}$, we set up $p(p-1)/2$ hypotheses: $\mathrm{H}_{\mathrm{null},ij} : \ Y_{ic} \perp\!\!\!\perp Y_{jc} \mid s_c$, for all $i, j \in \mathcal{V}$ and $i < j$. If $\mathrm{H}_{\mathrm{null},ij}$ is rejected, then $(i, j) \in \mathcal{E}$.

Although under the null, $S_{ij}$ asymptotically follows $N(0, 1)$, in practice, the null distribution of $S_{ij}$ may slightly deviate from $N(0, 1)$ due to unobserved factors. Existing multiple testing literature recommends estimating the mean and variance under the null, which has been proven to enhance testing result accuracy[11,12]. Consequently, we adhere to this recommendation and estimate the null mean $\hat{\mu}_0$ and variance $\hat{\sigma}_0^2$ by the method mentioned in Jin and Cai[13].

After deriving $\hat{\mu}_0$ and $\hat{\sigma}_0^2$, we employ the following FDR control procedure to designate edges. We reject $\mathrm{H}_{\mathrm{null},ij}$ if $|S_{ij} - \hat{\mu}_0|/\hat{\sigma}_0 \geq \hat{b}$, where

$$\hat{b} = \inf \left[ 0 \leq b \leq b_p : \frac{p(p-1)[1 - \Phi(b)]}{\max\{1, \sum_{1 \leq i < j \leq p} I[|S_{ij} - \hat{\mu}_0|/\hat{\sigma}_0 \geq b]\}} \leq \alpha \right],$$

$b_p = 2\sqrt{\log(N \vee p)}$, $\Phi$ is the CDF of $N(0, 1)$, and $\alpha$ is the FDR level, which is often set at 5%. If such $\hat{b}$ does not exist, set $\hat{b} = b_p$. The FDR control method is similar to what introduced in Xie and Li[8].

#### Criteria to determining problematic low-count genes.

We need to remove problematic low-count genes from the co-expression graph to avoid false positives. These genes are those satisfying all four criteria.

- Gene $i$ has a median read count $\leq 2$.
- The number of $S_{ij}$ passed the multiple testing threshold is greater than 10.
- Among all $S_{ij}$ passed the multiple testing threshold, the positive ones take up more than 90%.
- Among all the genes in $\widetilde{\mathcal{N}}_i^+ = \{j : S_{ij} > 0, (i, j) \in \mathcal{E}\}$, those with the same median as gene $i$ take up more than 90%.

#### Determining default connectivity cutoffs for identifying feature genes

We chose default cutoff parameters for co-expression network topology patterns by checking the distribution of $C_{1i}$, $C_{2i}$ and $C_{3i}$ of feature genes, pathway genes and hub genes in the Monoclonal dataset. The feature genes were collected from the GSEA MsigDB C2 curated human gene sets v2023.1[3–5,14–18] and the Uniprot database[19]. See "Experimental Data: Monoclonal" in Methods for details. The pathway genes and hub genes were collected in the following way.

- The pathway gene set were collected from the "KEGG_GLYCOLYSIS_GLUCONEOGENESIS", "KEGG_CITRATE_CYCLE_TCA_CYCLE", and "KEGG_OXIDATIVE_PHOSPHORYLATION" gene sets in GSEA MSigDB C2 curated human gene sets v2023.1. They include 144 cellular respiration related genes, which are functional in the same pathway but are not differentially expressed feature genes.
- The hub genes were collected in 3 studies on hub genes in colorectal cancer[20–22].

Human genes were converted to their mouse equivalents using `biomaRt` R package[23,24].

The distributions of $C_{1i}$, $C_{2i}$ and $C_{3i}$ in these three gene sets are shown in Note Fig. 3. Based on their distributions, by default, we require $C_{1i} > 5/(p-1)$ to exclude genes that does not show co-expression patterns with other genes. We require $C_{2i} > 0.4$ to filter out hub genes and $C_{3i} > 0.3$ to filter out pathway genes. Using these thresholds, we would be able to separate some feature genes from non-feature genes.

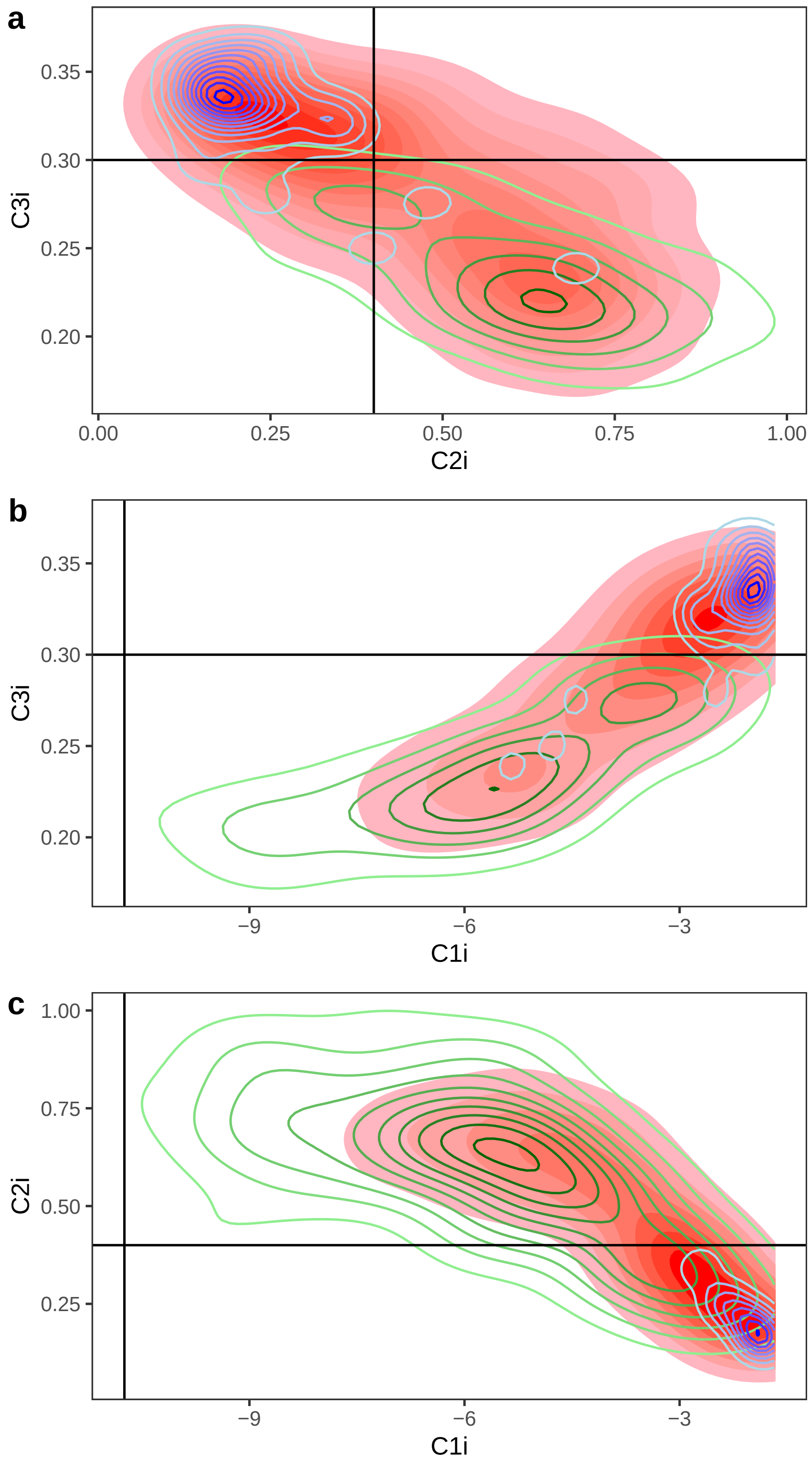


Note Figure 3: **Distributions of** $(log2C_{1,i}, C_{2,i}, C_{3i})$ **for feature genes, pathway genes, and hub genes**. **(a)-(c)** The 2D density contour plot of (a) $C_{2i}$ v.s. $C_{3i}$, (b) log2 $C_{1i}$ v.s. $C_{3i}$ and (c) log2 $C_{1i}$ v.s. $C_{2i}$ for feature genes (red), pathway genes (green) and hub genes (blue) in Monoclonal dataset. Darker color implies higher density. The black lines mark the default connectivity cutoffs.

### Supplementary Note 4: Simulation settings

#### Simulation settings of SD1

SD1 simulated 100 scRNA-seq datasets under identical settings. Each dataset consists of 6000 cells and 5000 genes. The cells are categorized into three cell types: 100 cells (1.7%) of type 1, 4900 cells (81.7%) of type 2, and 1000 cells (16.7%) of type 3. There are a total of 120 feature genes, with each cell type characterized by 50 feature genes. Notably, the feature genes of cell type 3 differ from those of cell type 1 and 2, while cell types 1 and 2 share 30 (60.0%) of their feature genes. The feature genes are simulated by applying fold changes to their baseline mean expression levels. All the feature genes are up-regulated, with fold changes greater than 1 in the corresponding cell type. In SD1, the rare cell type 1 can be considered to stem from non-rare cell type 2.

#### Simulation settings of SD2

SD2 comprises five cell types, each consisting of 1200 cells. Cell types 1, 3, and 5 are ending cell types, representing three distinct stages in the developmental process. Cell types 2 and 4 are transitional cell types between 1 and 3, and 3 and 5, respectively. The developmental lineage of cell types 1-2-3-4-5 follows the pattern: ending-transition-ending-transition-ending.

Within SD2, there are 220 feature genes among the 5000 genes. Each cell type exhibits 60 feature genes. Transitional cell types share 20 feature genes with their adjacent ending cell types, while the non-adjacent cell types do not share their feature genes. These feature genes can be categorized into nine sets, labeled as G1 through G9, based on the cell subpopulations where the genes are differentially expressed (Note Fig. 4). G1 and G9 contain 40 genes, while the remaining gene sets contain 20 genes. Sets G1, G3, G5, G7, and G9 comprise genes differentially expressed in a single cell type. Other sets are shared by neighboring cell types. Similar to SD1, all the feature genes are simulated by applying fold changes (> 1) to their baseline mean expression levels.

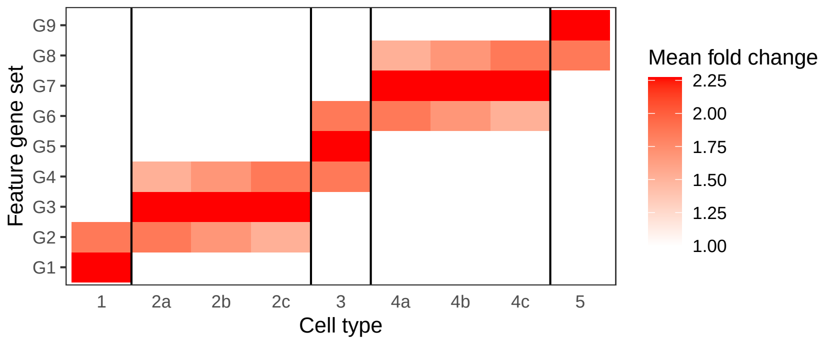


Note Figure 4: **Differential expression settings of SD2**. The fold changes of feature gene sets G1-G9 in different cell types. White color indicates a fold change equal to 1, suggesting that genes in the gene set are not differentially expressed feature genes in the specific cell type. Conversely, red color indicates the genes in the gene set are differentially expressed in the corresponding cell type, with darker shades of red representing larger mean fold change.

Each transitional cell type is further divided into three equal-sized cell subtypes (each cell subtype consists of 400 cells). These subtypes share the same feature genes but have varying expression level fold changes, mimicking the transitional process (Note Fig. 4). For example, cell type 2 is categorized into 2a, 2b, and 2c subtypes. Subtype 2a represents the transitional cells that are more similar to cell type 1 than cell type 3. In subtype 2a, the mean fold changes of G2 genes are the same as those in cell type 1 and mean fold changes of G4 genes are lower than those in cell type 3. Conversely, subtype 2c leans more toward cell type 3 than cell type 1. Subtype 2b falls between these extremes, with intermediate mean fold changes of G2 and G4 genes between those observed in cell types 1 and 3. Cell type 4 replicates the structure patterns in cell type 2.

[2] 10x Genomics. Pbmc from a healthy donor - granulocytes removed through cell sorting (10k), 2020. Single Cell Multiome ATAC + Gene Exp. Dataset by Cell Ranger ARC 1.0.0.
